# An Orc6 tether mediates ORC binding site switching during replication origin licensing

**DOI:** 10.1101/2025.05.09.652650

**Authors:** David Driscoll, Larry J. Friedman, Jeff Gelles, Stephen P Bell

## Abstract

During origin licensing, the origin recognition complex (ORC) loads two Mcm2-7 helicases onto DNA in a head-to-head conformation, establishing the foundation for subsequent bidirectional replication. Single-molecule experiments support a helicase-loading model in which one ORC loads both Mcm2-7 helicases at origins. For this to occur, ORC must release from its initial Mcm2-7 and DNA binding sites, flip over the helicase, and bind the opposite end of the Mcm2-7 complex and adjacent DNA to form the MO complex. Importantly, this binding-site transition occurs without ORC releasing into solution. Using a single-molecule FRET assay, we show that the N-terminal half of Orc6 tethers ORC to the N-terminal tier of Mcm2-7 (Mcm2-7N) during ORC’s binding-site transition. This interaction involves both the folded Orc6 N-terminal domain (Orc6N) and the adjacent unstructured linker and forms before ORC releases from its initial Mcm2-7 interaction. The absence of this interaction increases the rate of ORC release into solution, consistent with a tethering function. CDK phosphorylation of ORC inhibits the tethering interaction, providing a mechanism for the known CDK inhibition of MO complex formation. Interestingly, we identify mutations in the Orc6 linker region that support MO complex formation but prevent double-hexamer formation by inhibiting stable second Mcm2-7 recruitment. Our study provides a molecular explanation for a one-ORC mechanism of helicase loading and demonstrates that Orc6 is involved in multiple stages of origin licensing.

**Significance Statement:** Bidirectional DNA replication is critical for accurate and complete duplication of the genome. Eukaryotic organisms coordinate this through loading of two oppositely-oriented Mcm2-7 replicative helicases at origins of replication. Using single-molecule biochemical studies, we identified and characterized a tethering interaction during helicase loading that enables the helicase loader ORC (origin recognition complex) to flip between two Mcm2-7 and DNA binding sites to load the second helicases in the opposite orientation. This interaction is cell-cycle regulated as part of the mechanisms ensuring replication from a given origin initiates only once. Our findings have important implications for the multiple mechanisms of helicase loading and illustrate how single-molecule studies can complement structural studies to provide a full view of complex molecular assembly events.

## Introduction

Eukaryotic DNA replication begins at DNA sites known as origins of replication. During the G1 phase of the cell cycle, the process of origin licensing loads two copies of the Mcm2-7 replicative helicase onto origin DNA in an inactive, head-to-head conformation known as the double hexamer (Evrin et al., 2009; Noguchi et al., 2017; Remus et al., 2009). Upon S phase entry, these double hexamers are transformed into two active Cdc45-Mcm2-7-GINS (CMG) replicative helicases (reviewed in Bell and Labib 2016; Costa and Diffley 2022) that form the foundation for each replisome. Formation of double hexamers during G1 marks all potential origins of DNA replication and the orientation of the two Mcm2-7 helicases in the double hexamer ensures that subsequent initiation is bidirectional, a critical property ensuring complete genome replication.

Coordination of replication initiation events is critical to maintaining proper genomic content and ploidy. Loaded Mcm2-7 double hexamers require increased cyclin-dependent kinase (CDK) activity for CMG formation and activation. In *S. cerevisiae* cells, the same elevated CDK activity inhibits origin licensing outside of G1 phase by phosphorylating three helicase-loading proteins: Cdc6, Mcm2-7, and ORC (Nguyen et al., 2001). Phosphorylation of these proteins either lowers their effective concentration in the nucleus (Cdc6, Mcm2-7/Cdt1) (Drury et al., 2000; Tanaka and Diffley, 2002) or directly inhibits helicase loading (ORC) (Amasino et al., 2023; Chen and Bell, 2011; Frigola et al., 2013; Nguyen et al., 2001; Phizicky et al., 2018). By preventing origin licensing during S, G2, and M phases, this mechanism ensures that origins do not reinitiate during a single mitotic cell cycle thereby preventing genome re-replication.

Origins of replication in *S. cerevisiae* contain at least two DNA binding sites for the initiator protein, the origin recognition complex (ORC). Helicase loading begins with ORC binding to the highly conserved ARS consensus sequence (ACS) found at all origins (Figure S1A, step i, Bell and Stillman 1992; Broach et al. 1983; Zhang et al. 2023). In addition to the ACS, yeast origins include at least one, oppositely-oriented, weaker ORC DNA binding site, referred to as the B2 element (Chang et al., 2011; Marahrens and Stillman, 1992; Palzkill and Newlon, 1988; Wilmes and Bell, 2002). Importantly, the presence of two oppositely-oriented ORC binding sites is required for origin function (Coster and Diffley, 2017; Marahrens and Stillman, 1992). This juxtaposition of ORC binding sites enables ORC to load two Mcm2-7 helicases in the characteristic head-to-head orientation required for bidirectional replication (Costa and Diffley, 2022). In addition to these two DNA binding sites, analysis of the events of helicase loading have revealed that ORC has two distinct and important binding sites on Mcm2-7 (Figure S1A). After binding the ACS, ORC recruits Cdc6 followed by a complex between the Mcm2-7 helicase and Cdt1, forming the short-lived ORC-Cdc6-Cdt1-Mcm2-7 (OCCM) complex (Figure S1A, steps ii-iii, Ticau et al. 2015; Yuan et al. 2017). This complex involves ORC binding Mcm2-7 via extensive interactions between the C-terminal tiers of ORC and Mcm2-7, which we will refer to as the ‘OM interaction.’ The OCCM is lost through the sequential release of Cdc6 then Cdt1 (steps iv-v). Importantly, the loss of Cdt1 coincides with disruption of the OM interaction (Gupta et al. 2021). Rapidly after this event, ORC binds a second distinct region of this first Mcm2-7 to form the MO complex (step vi, Miller et al. 2019; Gupta et al. 2021). In contrast to the C-tier interactions involved in initial Mcm2-7 recruitment and the OCCM, ORC is bound to the opposite N-terminal tier of the Mcm2-7 helicase in the MO complex. These ‘MO interactions’ are primarily mediated by the smallest ORC subunit, Orc6 (Gupta et al., 2021; Miller et al., 2019). Importantly, in the MO complex ORC also binds to the oppositely-oriented B2 site on the DNA. The MO complex is required to recruit a second Cdc6 and Mcm2-7-Cdt1 complex, resulting in loading of the second Mcm2-7 in the opposite direction relative to the first followed by double-hexamer formation (steps vii-x, Champasa et al. 2019; Noguchi et al. 2017; Miller et al. 2019; Gupta et al. 2021). Although structural studies have provided critical information concerning the interactions between ORC and Mcm2-7 in the OCCM and MO complexes, the events that occur during the transition between these states are not fully understood.

Recent single-molecule studies showed that one ORC molecule is sufficient to load both Mcm2-7 complexes in the final double hexamer (Gupta et al., 2021). This observation means that ORC must change both its DNA and Mcm2-7 binding sites as it transitions between the OCCM and MO complexes. Prior studies have revealed that the ordered release of Cdc6 and Cdt1 from the OCCM coordinate ORC release from its initial DNA and Mcm2-7 binding sites. After Cdc6 release, ORC sliding on DNA releases it from the ACS DNA binding site (Figure S1A, steps iv-v, (Zhang et al., 2023). Simultaneous with Cdt1 release, the OM interaction between ORC and the C-terminal tier of Mcm2-7 is broken (Figure S1A, step v, Gupta et al. 2021). It remains unclear, however, what mechanism prevents ORC from releasing into solution as it fully releases from the DNA and flips its orientation to form new interactions with the N-terminal tier of Mcm2-7 and the oppositely-oriented B2 DNA element to form the MO-complex (Figure S1A, step vi).

The characteristics of the Orc6 subunit suggest a potential mechanism to retain ORC during its binding-site transition. The Orc1-5 subunits form a partial ring around double-stranded DNA at the ACS, however, Orc6 binds the periphery of this ORC ring (Li et al., 2018). In *S. cerevisiae*, Orc6 contains 3 domains: a C-terminal alpha helix that mediates binding to Orc3, a C-terminal TFIIB domain (Orc6C) that interacts with DNA, and a second N-terminal TFIIB domain (Orc6N) that is connected to the rest of Orc6 through a long, unstructured linker (Bleichert et al., 2013; Miller et al., 2019; Schmidt et al., 2022; Yuan et al., 2017). Consistent with this connection, the Orc6N domain is unresolved in many structural studies of helicase-loading intermediates, including DNA-bound ORC and the OCCM complex (Li et al., 2018; Yuan et al., 2017). In contrast, in the MO complex structure the Orc6N domain is resolved and interacts with the N-terminal Mcm2 A-domain (Mcm2N, Miller et al. 2019). This interaction along with the Orc6C domain interacting with the Mcm5 A-domain mediates ORC interaction with Mcm2-7 in the MO complex. Single-molecule studies showed the Orc6C domain interacts with the Mcm2-7 N-tier only after Cdt1 release (Gupta et al., 2021). In contrast, the long linker between the Orc6C and Orc6N domains raises the possibility that Orc6N could bind to the Mcm2-7 N-terminal domain prior to MO complex formation. Such an interaction would allow Orc6 to tether the rest of ORC to Mcm2-7 during its binding-site transition.

To understand the mechanism of the ORC binding-site switch during helicase loading, we combined colocalization single-molecule spectroscopy (CoSMoS, Friedman et al., 2006; Friedman and Gelles, 2015) with single-molecule Förster resonance energy transfer (smFRET) to monitor Orc6-Mcm2-7 interactions in real time. We show that ORC forms an interaction between its Orc6N domain and the Mcm2-7 N-terminal tier (Mcm2-7N) before the rest of ORC changes binding sites. This interaction depends on both the Orc6N domain and the Orc6 flexible linker and dramatically increases ORC retention after Cdt1 release. CDK phosphorylation of ORC inhibits formation and stability of this Orc6N-Mcm2-7N interaction. Mutations in the Orc6 linker also prevent stable second Mcm2-7 recruitment, demonstrating a second role for Orc6 in helicase loading. These studies reveal multiple roles for Orc6 during helicase loading and have important implications for the one-versus two-ORC pathways to form the Mcm2-7 double hexamer.

## Results

### Orc6N interacts with Mcm2-7N rapidly upon Mcm2-7 recruitment

A single ORC molecule transitioning between the OCCM and MO complexes could be explained by the Orc6N domain binding to Mcm2-7N, and the Orc6 flexible linker tethering the rest of ORC to Mcm2-7. If Orc6 forms such a tether before the ORC binding-site transition, then Orc6N should bind to Mcm2-7N before MO complex formation. To determine when Orc6N associates with the Mcm2-7N, we developed a smFRET assay for this interaction using *S. cerevisiae* proteins (Figure 1A, B). Orc6 was labeled at a peptide inserted adjacent to the N-terminal folded domain (position 107) using the SFP enzyme (ORC^6N-550^, Figure S1B, Zhou et al. 2007). The Mcm2-7 helicase was labeled at its N-tier by deleting the Mcm6 N-terminal extension (amino acids 1 to 103) and attaching a fluorescently-labeled peptide adjacent to this site using Sortase (Mcm2-7^6N-650^) (Figure S1B). Previous studies showed the Mcm6 extension is dispensable for helicase loading (Champasa et al., 2019). Importantly, bulk helicase-loading remained robust in reactions using one or both of these modified proteins (Figure S1C). Using total internal reflection fluorescence microscopy, we monitored the colocalization of ORC^6N-550^ and Mcm2-7^6N-650^ with individual origin DNA molecules (Friedman and Gelles, 2015, Figure 1A). Alternate excitation of the donor and acceptor fluorophores allowed observation of the association of either ORC or Mcm2-7 with DNA. Importantly, interactions between Orc6N and Mcm2-7N were monitored by determining the apparent FRET efficiency (*E*_FRET_) during donor excitation (Figure 1B). We restricted our analysis to events in which ORC and Mcm2-7 were both simultaneously present on DNA for at least 5 frames (>∼12s) to focus on productive ORC-Mcm2-7 interactions. We will refer to association events that meet these criteria as “stably recruited” Mcm2-7 helicases.

**Figure 1:**
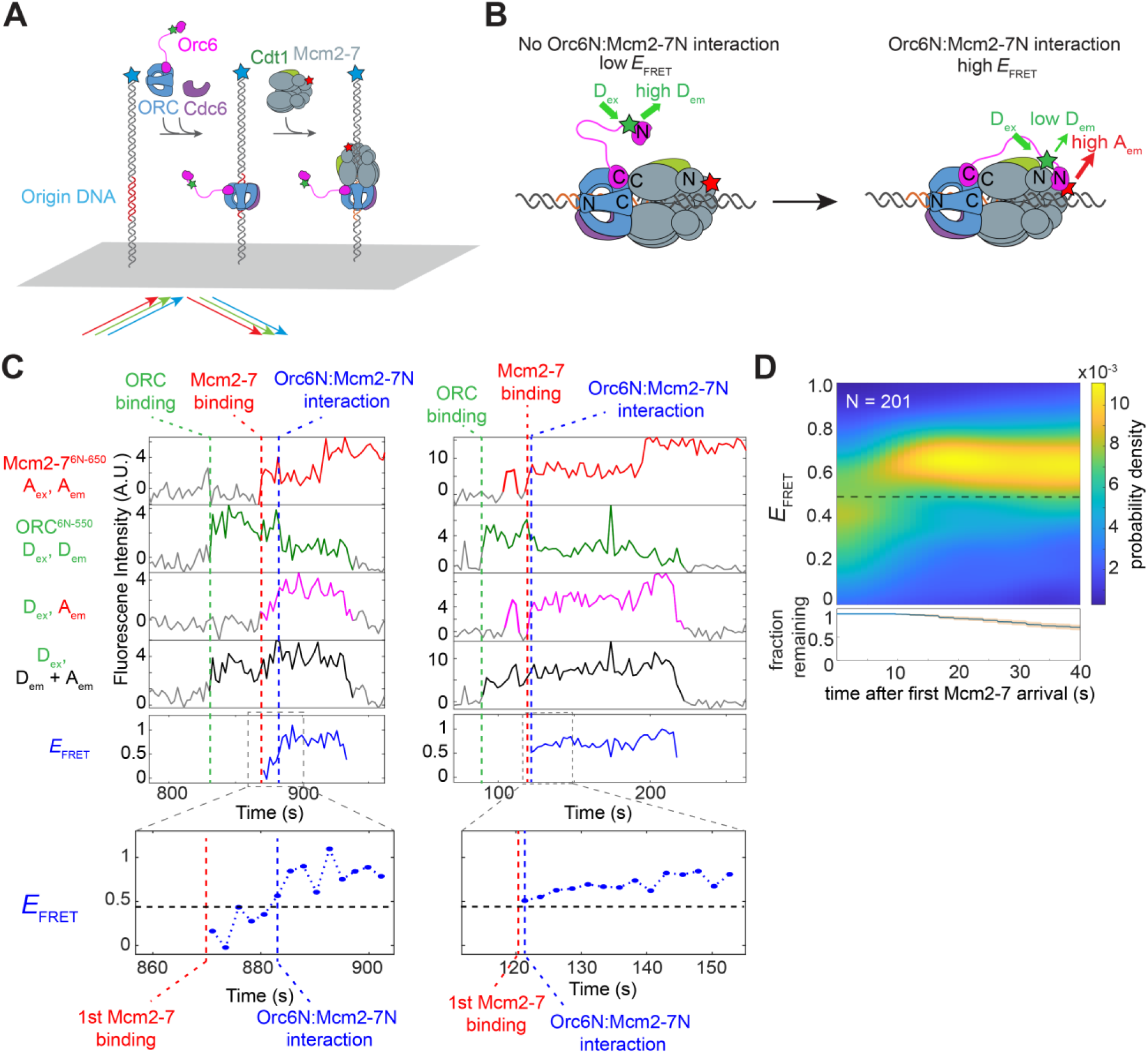
Orc6N interacts specifically with Mcm2-7N rapidly upon Mcm2-7 recruitment. A: Schematic of the single-molecule colocalization/FRET assay used in this study. DNA molecules containing an origin of replication were labeled with a blue-excited fluorophore and attached to a microscope slide. A solution containing all four helicase-loading proteins (with the indicated proteins labeled, stars) was incubated with DNA. Colocalization of green (donor) or red (acceptor) fluorophores with blue fluorophores was used as a proxy for protein-DNA associations (Friedman and Gelles, 2015). B: Detection of Orc6N:Mcm2-7N interaction by single-molecule FRET. In the experiment shown (A), ORC is labeled with a donor fluorophore (green star) adjacent to its N-terminal domain (ORC^6N-550^) and Mcm2-7 is labeled with an acceptor fluorophore (red star) at the N-terminus of Mcm6 (Mcm2-7^6N-650^). When both labeled proteins are present on a DNA (A, right), a low *E*_FRET_ is expected when Orc6N and Mcm2-7N do not directly interact (left) and elevated *E*_FRET_ is expected when Orc6N and Mcm2-7N interact (right panel). C: Representative traces showing ORC^6N-550^ and Mcm2-7^6N-650^ DNA associations and Orc6N:Mcm2-7N interactions. Top plot: acceptor emission during acceptor excitation (red, A_ex_, A_em_). Red dashed line indicates time of Mcm2-7^6N-650^ recruitment to DNA. Second plot: Donor emission during donor excitation (green, D_ex_, D_em_). Green dashed line indicates time of ORC^6N-550^ binding to DNA. Third plot: FRET, acceptor emission during donor excitation (pink, D_ex_, A_em_). Fourth plot: total emission during donor excitation (black, D_ex_, (D_em_ + A_em_)). Bottom plot, effective FRET efficiency (*E*_FRET_)(blue, D_ex_, A_em_/(D_em_ + A_em_)). Blue dashed line indicates the transition from a low *E*_FRET_ state to a high *E*_FRET_ state corresponding to Orc6N:Mcm2-7N interactions. Gray line segments represent background signal when no fluorescent protein is colocalized with the DNA molecule. Line segments with signal above background are colored and indicate bound protein. Inset highlights *E*_FRET_ values (blue) after Mcm2-7 arrival (red dashed line). Horizontal black dashed line inside inset represents the Orc6N:Mcm2-7N *E*_FRET_ threshold value (see Figure S2 and Methods for details). Blue dashed vertical line in inset represents time at which *E*_FRET_ values move from below to above threshold value. D: *E*_FRET_ distribution heat map for 201 DNA molecules where ORC^6N-550^ bound and recruited Mcm2-7^6N-650^. Time is after Mcm2-7^6N-650^ arrival. The *E*_FRET_ probability density (color scale) was calculated using only the DNA molecules with bound ORC^6N-550^ and Mcm2-7^6N-650^ that remained at each time point; the fraction of remaining complexes (blue curve) and 95% CI (orange shading) are shown in the bottom plot. Dashed line is the threshold value used to distinguish between low and high *E*_FRET_ states (see Figure S2 and Methods for details).

When both ORC^6N-550^ and Mcm2-7^6N-650^ colocalized with DNA, *E*_FRET_ values frequently and rapidly increased after Mcm2-7^6N-650^ recruitment (Figure 1C, S1D). We found that 91.5% (184/201) of stably recruited Mcm2-7^6N-650^ led to establishment of a high *E*_FRET_ state (defined as two consecutive *E*_FRET_ measurements above a defined threshold, Figure S2A). A heat map of the *E*_FRET_ distribution versus the time after Mcm2-7 recruitment revealed the timing of the *E*_FRET_ transition (Figure 1D). Upon first Mcm2-7 recruitment, the majority of molecules are in a low *E*_FRET_ state (peak at 0.33), but transition to a high *E*_FRET_ state (peak at 0.71) shortly afterwards (Figure 1D and S2A). Kinetic analysis showed a 7.2 ± 1.2 s (median ± S.E.) time to form the high *E*_FRET_ state after Mcm2-7-Cdt1 recruitment (Figure S1E), substantially faster than the 38.3 ± 2.8 s estimated median time of MO formation after Mcm2-7-Cdt1 recruitment (Gupta et al. 2021). Once formed, the median duration of the Orc6N-Mcm2-7N interaction was 26.4 ± 2.4 s, with a subset of the events showing much longer interactions (Figure S1F). The high *E*_FRET_ state, once formed, was almost always detected until ORC release. Indeed, in cases of successful recruitment of a second Mcm2-7 (Figure 1C and S1D), the end of the Orc6N-Mcm2-7N interaction typically occurred when ORC was released from the DNA. Because previous studies showed that the MO complex dissolves before ORC release (Gupta et al., 2021), we conclude that the Orc6N-Mcm2-7N interaction forms before and is lost after the MO complex ends.

Because Orc6N is tethered to the rest of ORC via an unstructured linker, we considered the possibility that the high *E*_FRET_ we observed reflected the general proximity of Orc6N to Mcm2-7 rather than a specific interaction with the Mcm2-7 N-tier. To distinguish between these possibilities, we performed the same analysis except using Mcm2-7 labeled at its C-terminal tier via Mcm2 (Mcm2-7^2C-650^, Figure S3A and S3B). Using Mcm2-7^2C-650^ and ORC labeled at the same site on Orc6 as above, we did not observe stable formation of a high *E*_FRET_ state (Figure S3C). Instead, a small drop in *E*_FRET_ values occurred at a similar time (∼8 sec) as the transition to the high *E*_FRET_ state seen when Mcm2-7 was labeled at its N-terminal domain (Figure S3D). This finding is consistent with binding of Orc6N to the opposite end of the helicase reducing an already weak *E*_FRET_ signal. We conclude that the elevated *E*_FRET_ observed in the FRET assay (Figure 1B) reflects a stable interaction between the Orc6N and Mcm2-7 domains. For reasons that will become clear, we subsequently refer to this high-FRET state (Figure 1D) as the Orc6 tether interaction and the assay presented in Figure 1 as the Orc6-tether assay.

### Two regions of Orc6 contribute to the tether interaction

Given the known interaction between the Orc6N and Mcm2N domains in the MO complex (Gupta et al., 2021; Miller et al., 2019), we asked if Orc6N is involved in the interaction observed in our FRET assay. To this end, we performed the Orc6-tether assay using ORC that lacked Orc6N (ORC^6ΔN-550^ in Figure S4A; Figure 2A, left). Interestingly, upon stable recruitment of Mcm2-7^6N-650^ by this ORC complex, we still observe a clear transition to a higher *E*_FRET_ state (Figure 2B, left, and Figure S5A). The timing and stability of the high *E*_FRET_ state measured when using ORC^6ΔN-550^ were largely unaltered (Figure 2C-D), however, one notable kinetic difference was observed. ORC^6ΔN-550^ showed a clear reduction in longer-lived tether interactions compared to ORC^6N-550^ (Figure 2D, >40s), consistent with the known defect in MO complex formation for this mutant (Gupta et al., 2021; Miller et al., 2019). In addition, there was a modest reduction in the percentage of molecules that reached the high *E*_FRET_ state (82.3% (102/124) for ORC^6ΔN-550^ vs. 91.5% for ORC^6N-550^ (184/201), Figure 2E), and the average high *E*_FRET_ value was reduced (high *E*_FRET_ state centered around 0.62 for ORC^6ΔN-550^ vs. 0.71 for ORC^6N-550^ (Figure S2A and S2B, left). Together, these findings suggest that, although the Orc6N domain is involved in the tether interaction, additional regions of Orc6 are likely to contribute to the interaction.

**Figure 2:**
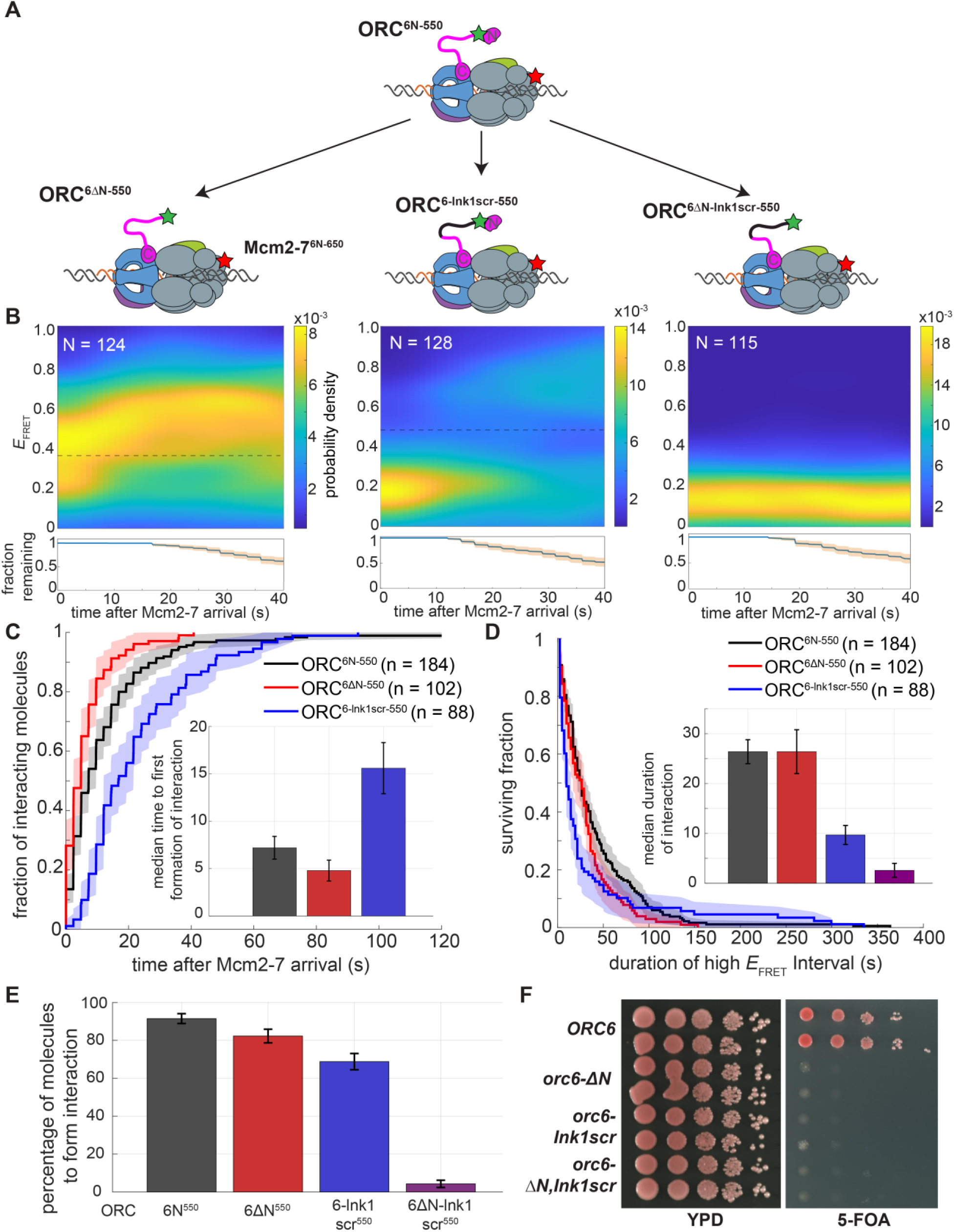
Orc6N and the adjacent linker region facilitate the Orc6 tether interaction. A: Schematic diagram of the Orc6 mutants used in these experiments. All mutants were labeled with a donor fluorophore (green stars) and used in the same helicase-loading assay shown in Figure 1 using Mcm2-7^6N-650^ labeled with an acceptor fluorophore (red stars). The black line represents the scrambled mutant of the lnk1 region of Orc6. See Figure S4A for mutant and labeling details. B: Heat maps showing the probability density of Orc6 tether interactions over time after Mcm2-7^6N-650^ arrival using the mutant constructs: ORC^6ΔN-550^ (left), ORC^6-lnk1scr-550^ (middle), and ORC^6ΔN-lnk1scr-550^ (right). Dashed line is the threshold value used to distinguish between low and high *E*_FRET_ states for each construct (see Figure S2B). Bottom plots show fraction of molecules remaining at the indicated times after first Mcm2-7 recruitment (orange shading shows 95% CI). C: Time to first formation of Orc6 tether interaction. Wild type (black), ORC^6ΔN^ (red), and ORC^6-lnk1scr^ (blue) are plotted relative to Mcm2-7^6N-650^ arrival. The Y-axis shows the fraction of molecules that formed Orc6 tether interaction. Shading represents 95% CI. To facilitate temporal comparison, molecules that did not reach a high *E*_FRET_ state were not included in the analysis. Inset: Bar graph showing median times (± S.E.) to reach high *E*_FRET_ state. D: Cumulative survival curve of initial Orc6 tether interactions. Wild type (black), ORC^6ΔN^ (red), and ORC^6-lnk1scr^ (blue) are plotted. Shading represents 95% CI. Only the first Orc6 tether interaction for a DNA molecule was considered for this analysis. The ORC^6ΔN-lnk1scr^ data was not plotted on the line graph for clarity due to few data points (n = 5). Molecules that never formed the tether interaction were eliminated from the analysis. Inset: Bar graph showing median durations (± S.E.) of high *E*_FRET_ state (purple bar is ORC^6ΔN-lnk1scr^ data). E: Percentage of molecules that formed the Orc6 tether interaction. The percentage (± S.E.) was determined by counting the number of ORC-Mcm2-7 interactions that reached the high *E*_FRET_ state (defined as two consecutive measurements above threshold) relative to total Mcm2-7 recruitment events. F: Both Orc6N and Orc6 lnk1 regions are essential. The phenotype of the *orc6* mutants used above was tested using a plasmid-shuffle assay (see Methods). Growth on 5-FOA selects against cells containing wild-type *ORC6* plasmid, revealing the functionality of the remaining mutant *orc6* allele. Ten-fold serial dilutions of cells were grown on the indicated media for 2 days at 30°C.

We next investigated the role of the unstructured Orc6 linker using the Orc6-tether assay. We focused on a linker region (aa 120-185) previously identified as important for viability (Orc6 lnk1, Chen et al. 2011). To eliminate effects due to altered linker length, we scrambled the Orc6 lnk1 region by altering the order but not the identity of its residues (Figure S4A, ORC^6-lnk1scr-550^). Like the WT Orc6 lnk1, the scrambled sequence is predicted to be unstructured (Figure S4B).

Scrambling the Orc6 lnk1 region had a significant impact on the characteristics of Orc6 tether interactions (Figure 2B, middle panel, Figure S5B). The median time for this mutant to form the interaction (i.e., the time to enter the high *E*_FRET_ state) is two-fold longer than WT ORC (Figure 2C, compare black to blue). Additionally, the median lifetime of the tether interactions with the ORC^6-lnk1scr-550^ mutant is more than two-fold shorter (Figure 2D). The latter finding is consistent with the frequent oscillations between high and low *E*_FRET_ observed in individual traces (Figure S5B) and the bimodal *E*_FRET_ values observed after ∼20 seconds (Figure 2B, middle panel). Consistent with a weaker interaction, ORC^6-lnk1scr-550^ showed a reduction in the percentage of molecules attaining the high *E*_FRET_ state (Figure 2E). These data suggest that the lnk1 region in Orc6 is important, but not solely responsible for the observed Orc6 tether interaction.

Because neither elimination of the Orc6N domain nor mutation of Orc6-lnk1 prevented the tether interaction, we tested the impact of combining these mutations (Figure 2A, ORC^6ΔN-lnk1scr-550^). This mutant results in a dramatic loss of the high *E*_FRET_ signal (Figure 2B, right panel, Figure S5C). Of the 115 instances where Mcm2-7^6N-650^ complexes are stably recruited by ORC^6ΔN-lnk1scr-550^, only 4.4% (5/115) transitioned to a high *E*_FRET_ state (compared to 91.5% of WT ORC^6N-550^ molecules, Figure 2E). In the rare instances that FRET is observed with ORC^6ΔN-lnk1scr-550^, the duration is much shorter than the WT or either single Orc6 mutant (Figure 2D inset). Together our data indicate that the Orc6N and Orc6-lnk1 regions work together to form the Orc6 tether interaction.

We also asked if these mutations have a phenotype *in vivo* (Figure 2F). Previous studies showed that deletion of Orc6N is lethal to cells (Chen and Bell, 2011). Using a plasmid shuffle assay, we tested whether the ORC^6-lnk1scr^ or ORC^6ΔN-lnk1scr^ are similarly defective. When any of these alleles are the only copy of Orc6 present *in vivo*, we see that the associated cells are inviable (Figure 2F). Thus, the Orc6N domain and the Orc6 lnk1 region are essential for Orc6 function.

### The Orc6 tether interaction retains ORC after Cdt1 release

For one ORC to load both Mcm2-7 helicases without leaving the site of helicase loading, ORC must remain linked to the site of loading throughout its binding-site transition. After release of Cdt1, OM interactions are broken and the MO complex is formed rapidly (median 8.3 ± 2.0 s, Gupta et al. 2021). The interaction between Orc6 and Mcm2-7N represents a possible mechanism to tether ORC to Mcm2-7 as it flips and changes binding sites. For such a mechanism to operate, the interaction would need to form before the OM interaction and the ORC-DNA interaction are disrupted and be retained during the transition to MO complex formation (Figure 3A).

**Figure 3:**
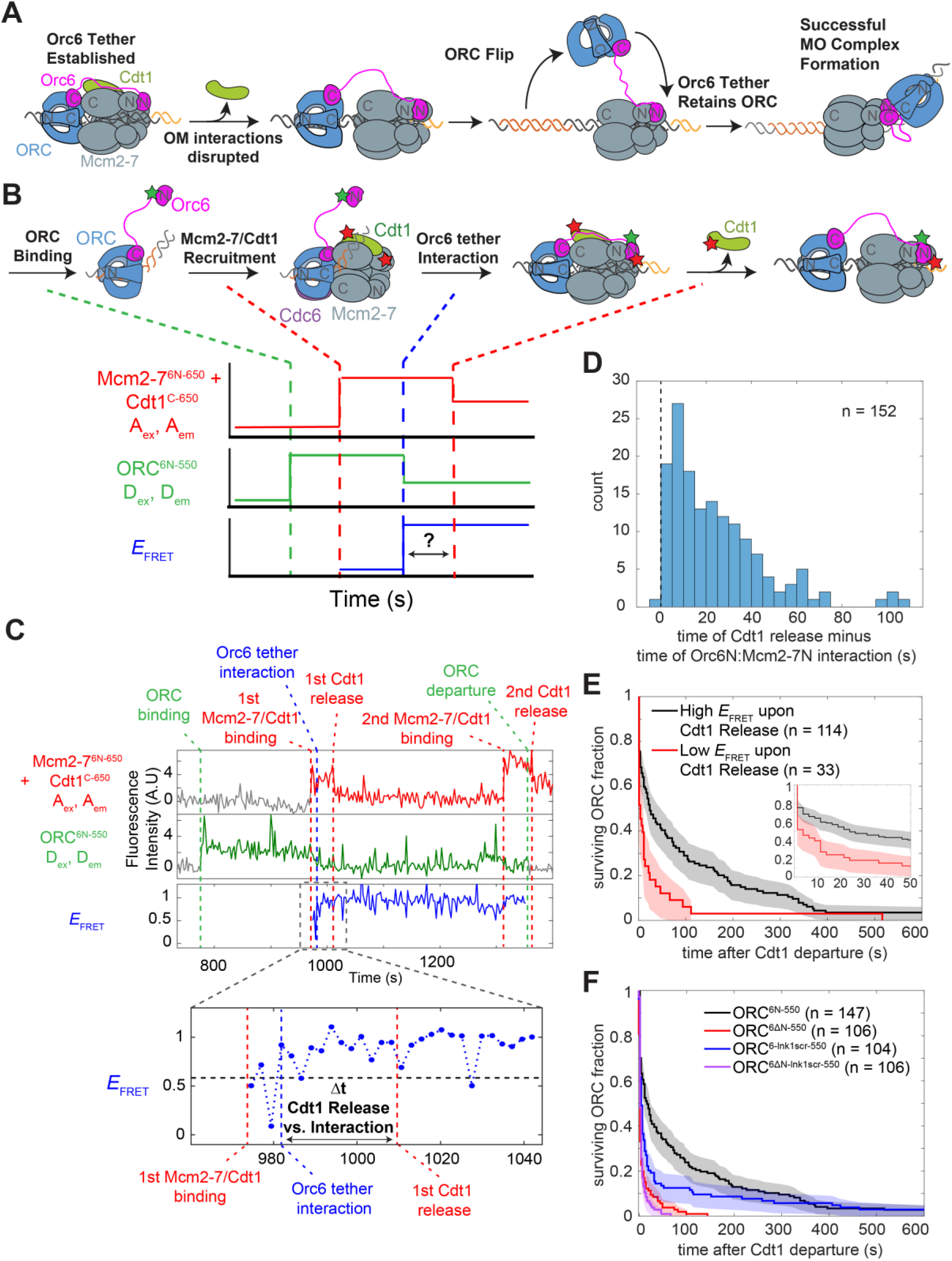
Orc6 interacts with Mcm2-7N before Cdt1 release and MO formation to tether ORC to Mcm2-7 during its binding site transition. A: Model for Orc6 tethering during OM to MO transition. Upon Cdt1 departure and breaking of OM interactions, Orc6 retains ORC at site of helicase loading to enable ORC flipping and formation of the MO complex. B: Experiment design for simultaneous assessment of Orc6 tether interaction and Cdt1 release. ORC^6N-550^ (donor) and Mcm2-7^6N-650^ (acceptor) are labeled as described previously (Figure 1). Cdt1^C-650^ is labeled with an additional acceptor fluorophore (red star). Diagram illustrates the expected sequence of species (top) and the single-DNA recording that would result if the tether interaction occurs before Cdt1 release (bottom). Binding of Mcm2-7^6N-650^-Cdt1^C-650^ causes an increase in red fluorescence. Orc6N and Mcm2-7N interaction results in a low to high *E*_FRET_ transition (blue plots). Cdt1 release causes a 50% decrease in red fluorescence with no change in *E*_FRET_ (top plot). The difference in the time of the latter two events (double-headed arrow) determines when Orc6N and Mcm2-7N interact relative to Cdt1 release. C: Representative single DNA molecule record from the experiment outlined in panel B. Traces are arranged as follows: Top plot: acceptor emission during acceptor excitation (red, A_ex_, A_em_). First red dashed line indicates time of first Mcm2-7^6N-650^-Cdt1^C-650^ recruitment to DNA, and second red dashed line indicates release of first Cdt1^C-650^. Third red dashed line indicates second Mcm2-7^6N-650^-Cdt1^C-650^ recruitment, and fourth red dashed line indicates second Cdt1^C-650^ release. Second plot: Donor emission during donor excitation (green, D_ex_, D_em_). First green dashed line indicates time of ORC^6N-550^ binding to DNA. Second green dashed line indicates time of ORC^6N-550^ release. Bottom plot, effective FRET efficiency (*E*_FRET_)(blue, D_ex_, A_em_/(D_em_ + A_em_)). Blue dashed line indicates the transition from a low *E*_FRET_ state to a high *E*_FRET_ state corresponding to Orc6 tether interactions. See Figure S6C for additional records. D: Histogram of times between Cdt1 release and the time of first Orc6N interaction with Mcm2-7N. Of 162 events that recruited both ORC^6N-550^ and Mcm2-7^6N-650^-Cdt1^C-650^, 152 events where a high *E*_FRET_ interaction was observed are shown. E: Survival plot showing fraction of ORC molecules retained after Cdt1 release. Data were segregated by whether ORC^6N-550^ and Mcm2-7^6N-650^ had a high (black) or low (red) *E*_FRET_ value at the time of Cdt1 release. DNA molecules where ORC was not detected at the time of Cdt1 release (15/162) were excluded from this analysis. Inset shows 0 to 50 seconds after Cdt1 release to highlight differences in early time points. F: Impact of Orc6 mutants on ORC retention after Cdt1 release. Survival plots are shown for WT ORC and ORC containing Orc6 mutations (black: WT ORC^6N-550^, red: ORC^6ΔN-550^, blue: ORC^6-lnk1scr-550^, purple: ORC^6ΔN-lnk1scr-550^). In contrast to E, the data are not separated by *E*_FRET_ state at time of Cdt1 release. DNA molecules where Mcm2-7 released before ORC (5/111 for ORC^6ΔN-550^, 10/116 for ORC^6ΔN-lnk1scr-550^) were excluded from this analysis.

Because the OM interaction ends when Cdt1 is released from Mcm2-7, we first investigated when this potential tethering interaction occurred relative to Cdt1 release. Although comparison with Cdt1 release data (Ticau et al., 2015) indicated that, on average, the Orc6N interacts with Mcm2-7N before Cdt1 release (Figure S6A), it was possible that this order only occurred in a fraction of loading events. To determine whether this order of events was consistently the case, we measured the times of these events at individual DNA molecules. To this end, we performed the Orc6-tether assay as in Figures 1 and 2 but included Cdt1 labeled with the same, red-excited fluorophore (Cdt1^C-650^) as Mcm6N (Figure 3B). When Mcm2-7^6N-650^-Cdt1^C-650^ is recruited to DNA, we see an initial increase in red-excited fluorescence corresponding to two red dyes (Figure 3B). Successful progression of the helicase-loading process results in the release of Cdt1^C-650^ while Mcm2-7^6N-650^ remains bound to DNA (Gupta et al., 2021; Ticau et al., 2015), which can be observed as a halving of red-excited fluorescence (Figure 3B). Importantly, the Cdt1^C-650^ used in this experiment does not exhibit strong *E*_FRET_ with the ORC^6N-550^ dye during helicase loading (Figure S6B).

Comparing the time of the first Cdt1 release to the time of the Orc6 interaction with Mcm2-7N revealed a clear order of events. When ORC^6N-550^ recruited Mcm2-7^6N-650^/Cdt1^C-650^ a high *E*_FRET_ state consistently formed (152/162, Figure 3C, S6C). Of the 152 recruitment events that exhibited high *E*_FRET_, all but one (151/152) showed that the Orc6 tether interaction formed before Cdt1 departure (median time 16.9 ± 1.7 s prior to Cdt1 release, Figure 3D). Thus, Orc6 consistently interacts with Mcm2-7N before Cdt1 release and MO complex formation.

We expect that lack of the Orc6 interaction with Mcm2-7N should decrease ORC retention on the DNA after the loss of the OM interaction. To test this prediction, we compared the times of ORC retention after Cdt1 release for events that did or did not exhibit the tether interaction at the time of Cdt1 release (Figure 3E). When the Orc6 tether interaction was present at the time of Cdt1 release, ORC was subsequently retained for a median time of 23.8 ± 8.8 s. In contrast, when the Orc6 tether interaction was absent at the time of Cdt1 release, ORC retention was much shorter (median time of 3.8 ± 3.4 s). Of the molecules that lacked the tether interaction at the time of Cdt1 release but exhibited longer ORC retention times (>10 s), more than half (5/9) formed the tether interaction within 10 s of Cdt1 release and this was true for all but one event that lasted more than 25 s.

We also asked how Orc6 mutants we tested in the tether assay impacted ORC DNA retention after Cdt1 release using the same assay. Both mutants that removed the Orc6 N-terminal domain showed strong defects in ORC retention (Figure 3F). In each case, ∼80% of molecules lacking Orc6N were released within 7.8 s (three frames) after Cdt1 release (ORC^6ΔN-550^, 80 ± 3.9% (85/106); ORC^6ΔNlnk1scr550^, 83 ± 3.6% (88/106)) compared to only 39 +/-4.0% (58/147) for WT ORC^6N-550^. Similarly, although 24 ± 3.5% (35/147) of WT ORC^6N-550^ were retained for more than 100 s on DNA, either very few (ORC^6ΔN-550^, 1 ± 0.9 % (1/106)) or none (ORC^6ΔNlnk1scr550^, 0% (0/106)) of the ORC complexes lacking Orc6N were retained for this period. These longer lasting events would include tethered ORC as well as ORC involved the MO complex and subsequent events. Mutating the linker region resulted in a hybrid set of defects. ORC^6-lnk1scr-550^ showed a significant increase in ORC molecules rapidly released after Cdt1 release relative to WT ORC^6N-550^ (63 ± 4.7% (66/104) released after 7.8 s). Unlike the mutants lacking Orc6N, mutation of the Orc6 linker alone showed an intermediate level of long-lasting ORC retention events, with 13 ± 3.2% (13/104) of these molecules being retained for longer than 100 seconds. These data suggest that the Orc6N domain is critical for retaining ORC for longer periods and that the Orc6 lnk1 region facilitates the establishment of these interactions (see Discussion). Together, our studies of ORC retention after Cdt1 release strongly support a model in which Orc6 establishes a tether interaction with Mcm2-7 before Cdt1 release that retains ORC at the site of helicase loading during its binding-site transition.

### ORC phosphorylation interferes with the Orc6 tether

CDK phosphorylation of ORC robustly inhibits helicase loading *in vivo* and *in vitro* (Amasino et al., 2023; Chen and Bell, 2011; Nguyen et al., 2001). Orc6 is one of two ORC subunits modified by CDK, and recent studies demonstrated that Orc6 phosphorylation fully inhibits MO-complex formation (Amasino et al., 2023). Interestingly, the CDK modification sites on Orc6 are either in or proximal to the Orc6-lnk1 sequence (Nguyen et al., 2001), raising the possibility that ORC phosphorylation interferes with formation of the Orc6 tether interaction. To address this hypothesis, we performed the Orc6-tether assay using CDK-phosphorylated ORC^6N-550^.

ORC phosphorylation (both Orc2 and Orc6) altered several characteristics of the tether interaction (Figure 4A and B). First, ORC phosphorylation delayed the median time of tether formation more than two-fold relative to unphosphorylated ORC (Figure 4C). More importantly, the lifetime of the phosphorylated ORC interaction is two and half times shorter than with unphosphorylated ORC (Figure 4D). We also observed more frequent transitions between high and low *E*_FRET_ states compared to unphosphorylated ORC^6N-550^ (Figure 4A, right). *E*_FRET_ values for phosphorylated ORC^6N-550^ split into a distinct high and low *E*_FRET_ states after Mcm2-7 recruitment with more data in the low *E*_FRET_ region (Figure 4B). Consistent with a defect in tether formation, only 53.9% (77/143) of the Mcm2-7^6N-650^ molecules recruited by phosphorylated ORC^6N-550^ ever exhibited a stable high *E*_FRET_ state, compared to 91.5% (184/201) for unphosphorylated ORC (Figure 4E).

**Figure 4:**
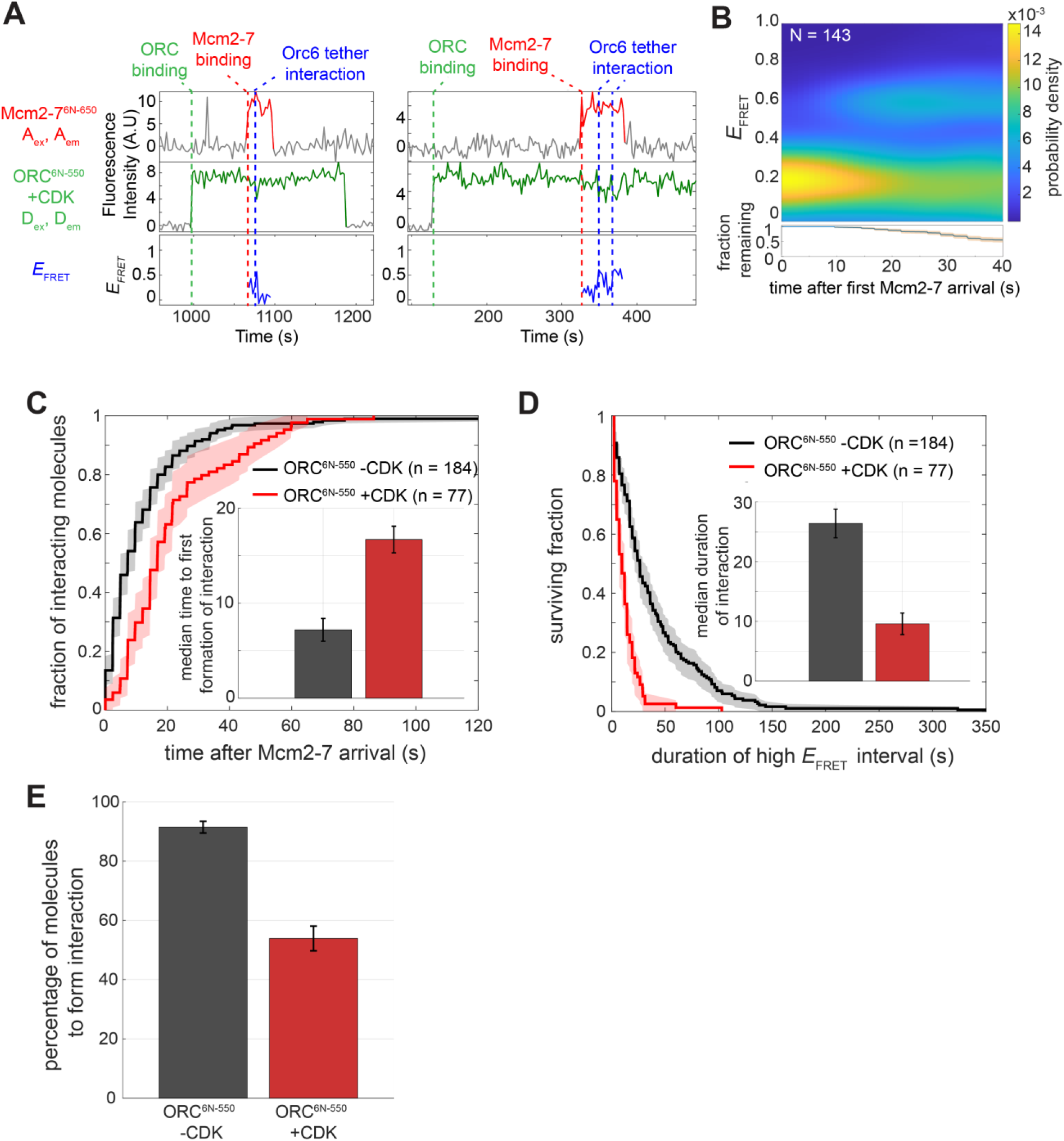
Phosphorylation of ORC inhibits the Orc6 tether interaction. A: Two representative single DNA molecule records of Orc6-tether assay using phosphorylated ORC^6N-550^, plotted as described in Figure 3C. B: Heat map showing the probability density of Orc6 tether interactions after Mcm2-7^6N-650^ arrival using ORC^6N-550^ phosphorylated by CDK. Lower plot shows the fraction of molecules remaining and 95% CI (blue curve, orange shading). C: Time to first formation of the Orc6 tether interaction. Unphosphorylated (black, same data as Figure S1E) and phosphorylated (red) ORC^6N-550^ are plotted relative to Mcm2-7^6N-650^ arrival. ORC was phosphorylated by CDK as described (Amasino et al., 2023). Shading represents 95% CI. To facilitate temporal comparison, molecules that never formed the tether interaction were eliminated from the analysis. Median time to first formation (± S.E.) of interaction for ORC^6N-550^ -CDK (black) and ORC^6N-550^ +CDK (red) (inset) was determined as in Figure 2C. D: Cumulative survival curve of initial Orc6 tether interactions for unmodified (ORC^6N-550^ – CDK, black) or CDK-modified ORC (ORC^6N-550^ +CDK, red). Shading represents 95% CI. Molecules that never formed the Orc6 tether interaction were eliminated from the analysis. Only the first interaction for a DNA molecule was considered for this analysis. Median durations of interaction (± S.E.) were determined as in Figure 2D. E: Percentage of molecules that formed the tether interaction are plotted. This percentage (± S.E.) was determined by counting the number of Orc6 tether interactions that reached the high *E*_FRET_ state compared to total Mcm2-7 recruitment events.

To address the contribution of Orc6 versus Orc2 phosphorylation, we repeated these experiments with ORC mutations that prevented phosphorylation of Orc6 (ORC^2phos-6N-550^) or Orc2 (ORC^6phos-6N-550^, Nguyen, Co, and Li 2001). We focused on the impact of each subunit’s phosphorylation on the time to formation and duration of the Orc6 tether interaction. Only Orc6 phosphorylation significantly delayed tether formation (Figure S7A-C). In contrast, phosphorylation of either Orc2 or Orc6 reduced the interaction lifetimes (Figure S7D-F).

CDK phosphorylation of Orc6 occurs in or proximal to the Orc6 lnk1 region (Chen et al., 2007; Nguyen et al., 2001). Given the overlapping targets, we compared the effects of Orc6 phosphorylation to the ORC^6-lnk1scr-550^ mutant. Interestingly, the distributions of initial times to tether formation are remarkably similar for the two ORC complexes (Figure S7G). The median lifetimes of the tether interactions obtained were also similar (12.0 ± 1.3 s vs. 9.7 ± 1.9 s, Figure S7F). However, when looking at the full distribution of interaction durations, we noticed a sub-population of ORC^6-lnk1scr-550^ mutant events that exhibited long-lived tether interactions that was absent when Orc6 was phosphorylated (Figure S7H). This observation raises the possibility that a subset of ORC^6-lnk1scr-550^ molecules can form one or more downstream complexes (e.g., MO complex) in helicase loading.

### Orc6 lnk1 is important for Mcm2-7 double-hexamer formation

To investigate the role of the Orc6 linker in the formation of complexes downstream of the OCCM, we performed a previously described single-molecule MO-complex-formation assay (Figure 5A, Gupta et al. 2021). Briefly, ORC was labeled with a donor fluorophore on the Orc6 C-terminus, and Mcm2-7 is labeled with an acceptor fluorophore on the Mcm3 N-terminus. When Mcm2-7 is initially recruited by ORC and forms the OCCM complex, these dyes are far apart and exhibit low *E*_FRET_. When ORC flips over Mcm2-7 to form the MO complex, the donor fluorophore on the Orc6 C-terminus is brought into close proximity with the acceptor fluorophore on the Mcm3 N-terminus resulting in a transition to a high *E*_FRET_ state (Figure 5A).

**Figure 5:**
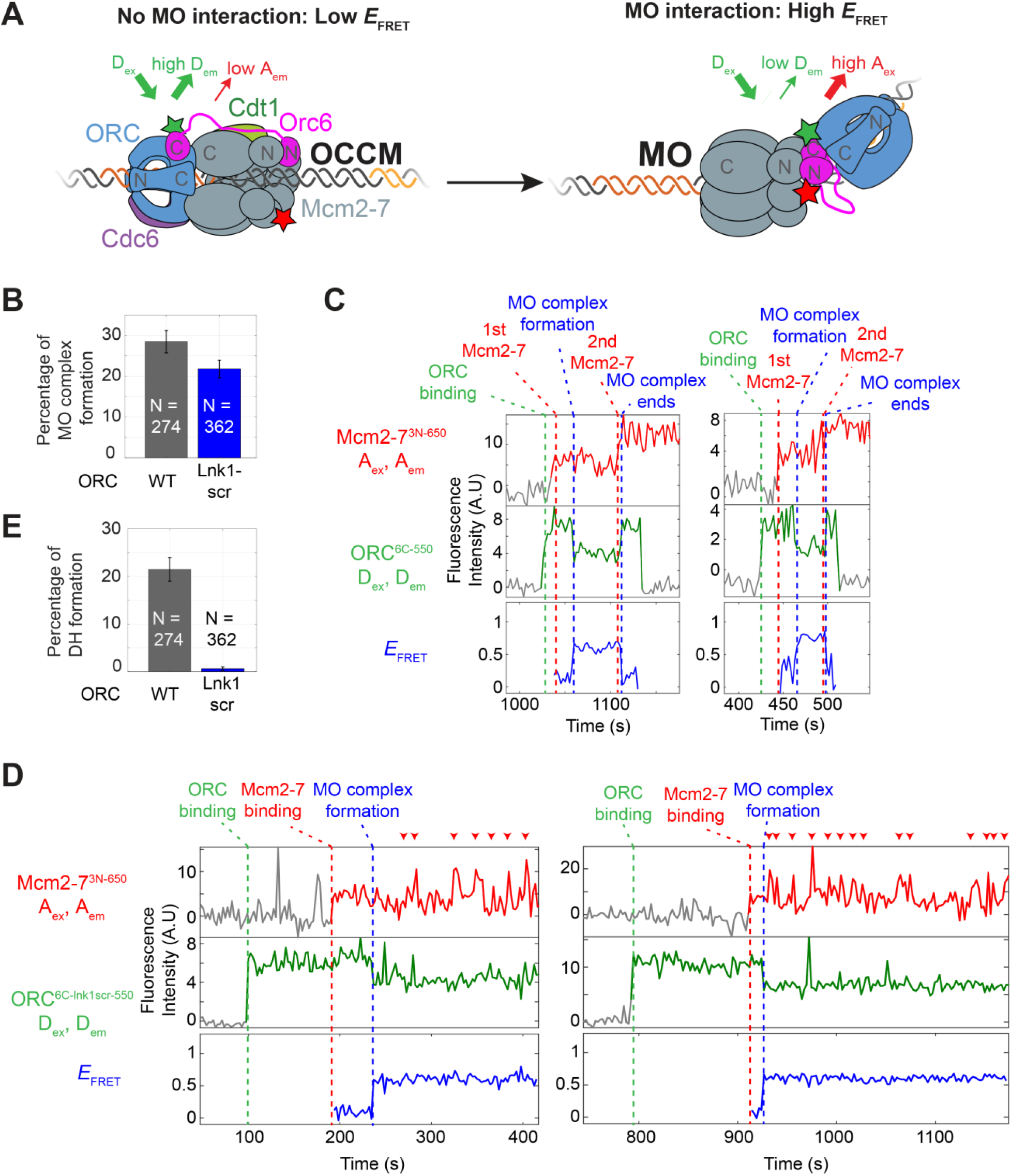
Orc6 lnk1 is important for stable second Mcm2-7 recruitment. A: Single-molecule FRET assay for MO complex formation. Reactions were performed as described (Gupta et al., 2021) using either WT (ORC^6C-550^) or ORC containing the Orc6 lnk1scr mutation (ORC^6C-lnk1scr-550^). ORC was labeled at Orc6 C-terminus (ORC^6C-550^) and Mcm2-7 was labeled at Mcm3 N-terminus (Mcm2-7^3N-650^, Gupta et al. 2021). B: Mutations in the Orc6 lnk1 region result in reduced MO complex formation. Bar graph showing the percentage (± S.E.) of 1^st^ Mcm2-7^3N-650^ binding events that resulted in formation of the MO complex for WT ORC^6C-550^ (black) and ORC^6C-lnk1scr-550^ (green). C: Representative traces of MO formation assay experiment using ORC^6C-550^. Plots are arranged as described in Figure 3C. D: Representative traces of MO formation assay experiment using ORC^6C-lnk1scr-550^. Plots are arranged as in Figure 5C. Red arrowheads indicate failed Mcm2-7 recruitment attempts. E: Mutations in the Orc6 lnk1 region cause severe defects in double-hexamer formation. Bar graph showing the percentage (± S.E.) of 1^st^ Mcm2-7^3N-650^ binding events that resulted in formation of an Mcm2-7 double hexamer for WT ORC^6C-550^ (black) and ORC^6C-lnk1scr-550^ (green).

Interestingly, although both ORC^6C-550^ and ORC^6C-lnk1scr-550^ showed interactions between Orc6C and Mcm2-7N, the mutant complex showed strong defects in downstream events. Performing the MO formation assay using ORC^6C-lnk1scr-550^ revealed a modest defect in the percentage of Mcm2-7 recruitment events that result in MO complex formation (Figure 5B). However, a striking phenotype was observed when later steps in the reaction were examined. For WT ORC^6C-550^, the MO complex dissociates shortly after 2^nd^ Mcm2-7 recruitment (Figure 5C, second vertical blue-dashed line). In contrast, the ORC^6C-lnk1scr-550^ mutant is trapped in the MO complex for extended periods of time as indicated by the persistent high *E*_FRET_ signal (Figure 5D). Although the MO complexes containing ORC^6C-lnk1scr-550^ repeatedly recruit 2^nd^ Mcm2-7 complexes, the recruited helicases are rapidly released (Figure 5D, red arrowheads). This results in a severe defect in DH formation compared to WT ORC (Figure 5E). Consistent with this defect, ensemble helicase loading assays using ORC^6C-lnk1scr-550^ show a strong defect in salt-stable Mcm2-7 helicase loading (Figure S8). These data indicate an important role for the Orc6 linker to stably recruit the second Mcm2-7 complex.

## Discussion

Previous studies showed that one ORC can mediate loading of both Mcm2-7 helicases at origins (Gupta et al., 2021; Ticau et al., 2015). This requires ORC to change its binding sites on both the first Mcm2-7 and DNA without being released into solution. In this study, we identified an interaction between the N-terminal half of Orc6 and the N-tier of Mcm2-7 that acts as a tether between ORC and Mcm2-7 during these binding site transitions. We showed that the interaction occurs with appropriate timing and functions to retain ORC during this transition. In addition, we identified a new function for the Orc6 lnk1 domain during recruitment of the second Mcm2-7.

### A tether function for Orc6 during helicase loading

Based on our findings, we propose a model in which Orc6 performs a tether function during helicase loading (Figure 6). Upon recruitment of Mcm2-7 to DNA-bound ORC-Cdc6, ORC embraces the first Mcm2-7 complex by first interacting with the Mcm2-7 C-terminal domains via the Orc1-5 C-tier along with Cdc6 (OM interactions). Shortly afterwards, the Orc6-N terminal and lnk1 regions localize to the Mcm2-7 N-terminal tier, forming a “tether” between ORC and the first Mcm2-7 (Figure 1). The interaction forms efficiently and rapidly once Mcm2-7 is recruited and is present well before Cdt1 release (Figure 3). Because Cdt1 release occurs simultaneously with the breaking of OM interactions and anticipates the formation of the MO complex (Gupta et al., 2021), this timing means the tether interaction is present prior to the ORC binding-site switch. In successful double-hexamer formation events, the Orc6 tether-associated FRET interactions last until after recruitment of a second Mcm2-7 complex (and therefore throughout MO complex formation) (Figures 1 and 3). This interaction keeps ORC bound to the site of helicase loading as it flips over the 1st Mcm2-7 helicase to form new interactions with the N-terminal domains of Mcm2-7 and the inverted B2 DNA binding site.

**Figure 6:**
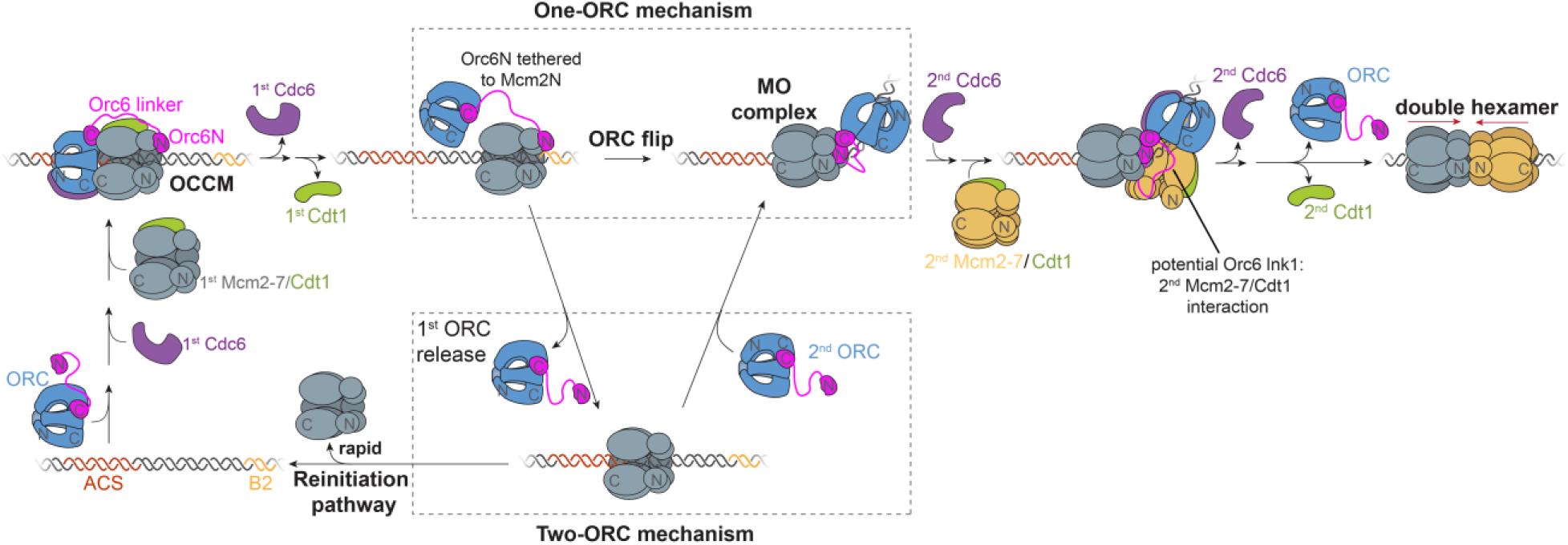
Model for Orc6 tether function during helicase loading. We propose that Orc6N interacts with Mcm2-7N rapidly after Mcm2-7 recruitment. This enables ORC to remain bound to the site of helicase loading when Cdt1 release occurs and OM interactions are broken, and tethers ORC to Mcm2-7 during its flip over Mcm2-7 to form the MO complex. See text for additional discussion.

The rapid and nearly quantitative formation of this tethering interaction upon first Mcm2-7 recruitment has interesting implications for the mechanism of helicase loading. These findings strongly suggest that in most cases (>90% in our experiments) the first approach taken to form a double hexamer is a single-ORC mechanism. Because the tether interaction forms between ORC and Mcm2-7 that are already bound to one another, an intramolecular event, even high concentrations of ORC would be unlikely to change this percentage. Multiple aspects of origin structure and helicase loading would prevent the involvement of a second ORC at this point. First, very few origins are arranged such that the ACS and B2 would be spaced appropriately to allow a second ORC to interact with the B2 element and the N-terminus of the first Mcm2-7 (Lim et al., 2024). For example, the B2 element at *ARS1* is in the central channel of the first Mcm2-7 upon initial recruitment, thus, preventing other proteins access to this DNA binding site until Mcm2-7 slides away from its initial position (Yuan et al., 2017; Zhang et al., 2023). Second, because the tethering interaction almost certainly involves the same regions of the Orc6 and Mcm2 involved in the MO interaction, the Orc6 tether interaction would interfere with a second ORC forming the MO complex.

Despite the likelihood that a one-ORC mechanism is the first approach used for origin licensing, this does not mean that it is the only mechanism used. Our single-molecule studies have also observed two-ORC reactions, albeit at a lower rate than the one-ORC mechanism (Gupta et al., 2021). A likely explanation for two-ORC events is the frequent release of ORC as it attempts to make its binding site transition. Even when a tether interaction was present as Cdt1 was released, ∼25% of ORC molecules released within one video frame (∼2.6 s) after Cdt1 release (28/114, Figure 3E inset). When the tether is not present as Cdt1 is released, approximately half of ORC molecules are released in the same time frame (16/33, Figure 3E inset). Therefore, we see ORC release at this stage as a common failure point in the one-ORC mechanism. The release of the first ORC provides an opportunity for a second ORC to form the MO complex and complete helicase loading (Figure 6, two-ORC mechanism). The time for a second ORC to form the MO complex is limited, however. If the first ORC fails to form the MO complex, it will also fail to hold the first Mcm2-7 in a stable closed-ring state (Amasino et al., 2023; Miller et al., 2019). A single Mcm2-7 alone on DNA is a relatively unstable state, having a median lifetime of 11.2 ± 2.0 s (Gupta et al., 2021). Release of the first Mcm2-7 prior to stabilization by binding a second ORC binding would reset the licensing process (Figure 6, reinitiation pathway). Thus, after first ORC release, a race between a second ORC binding and Mcm2-7 release would determine the extent of use of the two-ORC pathway. Thus, as ORC concentration increases so would the likelihood of a two-ORC pathway being used and it is very likely that both pathways are used in cells.

Why have electron microscopy studies of the OCCM complex structure (Yuan et al., 2017) not revealed an interaction between Orc6N and Mcm2-7N? The OCCM structure was trapped using a slowly hydrolysable ATP analog, ATPγS, to prevent Cdc6 departure and downstream steps (e.g., Mcm2-7 ATP hydrolysis which is required for subsequent translocation/sliding, Zhang et al. 2023; Butryn, Greiwe, and Costa 2025). In contrast, all reactions in the present study were performed with ATP. The discrepancy between the electron microscopy and FRET results suggests that there is an undetermined ATP-dependent step required to form the Orc6 tether interaction. We attempted to perform the tether assay with ATPγS, however, we were unable to obtain results due to poor association of ORC with DNA under these conditions. Although not detected by electron microscopy, crosslinking-mass spectrometry data from the same study (Yuan et al., 2017), also conducted in the presence of ATPγS, identified a crosslink between the Orc6 lnk1 sequence and the N-terminal A domain of Mcm2. Thus, the beginning of the interaction between Orc6N and Mcm2-7N may be occurring in the OCCM, although the longer-lasting interaction that we observe is not.

### Both the Orc6N domain and Orc6 lnk1 sequence contribute to Orc6 tether interaction

The data presented in this study reveal a role for both the Orc6N domain and the Orc6 lnk1 sequence in forming the Orc6 tether interaction. The Orc6N domain binds to Mcm2N in the MO structure (Miller et al., 2019) and deletion of the Orc6N domain prevents MO complex formation (Gupta et al., 2021; Miller et al., 2019). It is, therefore, most likely that the Orc6N domain interacts with the Mcm2-7N during tethering using the same or a similar interface. How the Orc6 linker contributes to the interaction observed between Orc6N and the first Mcm2-7N is less clear. This region of the Orc6 protein has not been resolved in any ORC-containing structures to date. Previous co-immunoprecipitation studies showed that the Orc6N + lnk1 (aa 1-185) region binds Cdt1 (Chen and Bell 2011), suggesting that Orc6 lnk1 contributes to tether interactions by binding Cdt1. Such an interaction raises the interesting possibility that Cdt1 release stimulates ORC reorientation during the flipping process by changing its interactions. Consistent with this idea, we found that once released from the ACS, ORC exhibits short ACS rebinding events only before Cdt1 release but never after (Zhang et al., 2023).

Although mutations in Orc6 lnk1 led to a longer time to form Orc6 tether interactions as well as shorter duration of these interactions (Figure 2C, D), these mutations did not eliminate long periods of ORC retention after Cdt1 release (Figure 3F) or MO FRET formation (Figure 5B). In contrast, deletion of Orc6N dramatically reduces ORC retention at the DNA after Cdt1 release (Figure 3F) and abolishes MO-complex formation (Gupta et al., 2021; Miller et al., 2019). These findings suggest that the Orc6 lnk1 interaction helps to position Orc6 N-terminus for tether formation but is not essential for tethering ORC during its binding site transition. This model is supported by the observation that, although mutation of Orc6 lnk1 leads to shorter interactions for the majority of ORC molecules, a subpopulation of Orc6 lnk1scr molecules form long-lived tether interactions (see bimodal curves in Figure 2D, 3F). A model in which Orc6 lnk1 interacts with Cdt1 to facilitate Orc6N interactions with Mcm2N in the first Mcm2-7, fits with the tether being mediated by Orc6N, as Cdt1 is no longer present during the ORC binding-site transition.

It has been proposed that the Orc2 IDR could have a role in tethering ORC to the site of helicase loading (Wu et al., 2024). This conclusion is based on the finding that the first 200 amino acids of Orc2 IDR interact with purified OCCM complexes (Wu et al., 2024). Another study found that Orc2 IDR residues 190-231 interact with Mcm5 and the Orc6 C-terminal domain in the MO complex (Lim et al., 2024). Because Orc6C is only proximal to this region of Mcm5 after ORC has flipped to the MO conformation, any tethering interaction involving Orc2 IDR would be with Mcm5 alone. This more limited interaction could explain the weak/transient nature of the Orc2 IDR-OCCM interactions (Wu et al., 2024). We show that deletion of Orc6N alone leads to rapid release of ORC after Cdt1 release (Figure 3F), suggesting that Orc2 IDR is not sufficient to retain ORC during the binding-site switch. We propose that a role of the Orc2 IDR in tethering could support but not substitute for the Orc6 tether interaction and likely plays a more prominent role in stabilizing the MO complex (Lim et al., 2024; Wu et al., 2024).

Interestingly, our findings also reveal a role for the Orc6 linker after MO complex formation. ORC can recruit the first Mcm2-7 and form an OCCM complex without Orc6 (Fernández-Cid et al. 2013). Although current models based on structural studies suggests that recruitment of the second Mcm2-7 helicase uses the same interactions as the first (Bell and Labib, 2016; Costa and Diffley, 2022), a role for Orc6 in stable recruitment of the second Mcm2-7 suggests otherwise. How could Orc6 be involved in recruitment of the second but not the first Mcm2-7? One possibility is that the ORC^6-lnk1scr^ mutant places the Orc6C domain in close proximity to the first Mcm2-7 N-tier but does not form a proper MO complex. Although we did not detect any difference in the high *E*_FRET_ states adopted by WT ORC^6N-550^ and ORC^6C-lnk1scr-550^ in our MO assay (Figure S8B), it is possible that alterations that impact second Mcm2-7 recruitment are not sufficient to change the *E*_FRET_ signal. Alternatively, the Orc6 lnk1 region may form an uncharacterized interaction with the second Mcm2-7-Cdt1 complex that facilitates its stable loading. Such an interaction could be related to the interaction we observe between the Orc6 lnk1 region and the first Mcm2-7-Cdt1 complex (ORC^6-lnk1scr-550^, Figure 2).

### CDK phosphorylation of ORC inhibits Orc6 tether interactions

Previous studies have shown that phosphorylation of ORC inhibits MO complex formation (Amasino et al., 2023; Lim et al., 2024). Our data suggests that this effect is at least partly due to inhibition of tether formation (Figure 4). Phosphorylation of Orc6 alone, but not Orc2 alone, decreases the rate of formation of this interaction (Figure S7). Based on this data, as well as the fact that Orc6 phosphorylation sites are either in or proximal to the Orc6 lnk1 sequence (Nguyen et al., 2001), we propose that Orc6 phosphorylation directly inhibits tether formation. Phosphorylation of either Orc6 or Orc2 reduces the duration of Orc6N-Mcm2-7N interactions (Figure S7). Again, the proximity of the Orc6 phosphorylation sites to Orc6N suggests that modification of Orc6 directly interferes with binding of the Orc6N to Mcm2-7N, reducing its duration. The mechanism of Orc2 phosphorylation reducing Orc6 tether duration is less clear but is likely related to interactions between the Orc2 IDR and Mcm2-7 in either the OCCM (Wu et al., 2024) or the MO complex (Lim et al., 2024; Wu et al., 2024) helping to stabilize ORC:Mcm2-7 interactions.

### Implications for origin licensing in other organisms

A major difference between replication initiation in budding yeast and most other eukaryotic organisms is the lack of specific sequences driving helicase loading in most organisms. The one-ORC mechanism seems particularly useful in a situation in which there are no high affinity ORC binding sites on DNA since this would reduce the need for recruitment of two ORC molecules to low-affinity binding sites. Consistent with a similar mechanism being used, recent studies of human origin licensing showed that a similar MO complex can be detected (Weissmann et al., 2024; Yang et al., 2024). Nevertheless, a key element of the one-ORC mechanism is changed. Orc6 in human cells (HsOrc6) is not robustly bound to the remaining ORC subunits and is not required for all helicase loading (although it improves the efficiency of the reaction) (Weissmann et al., 2024; Wells et al., 2025; Yang et al., 2024). In addition, the location and length of the linker is changed. In contrast to *S. cerevisiae* Orc6, HsOrc6 has the two TFIIB-related domains involved in MO complex formation located adjacent to one another at the N-terminus of the protein. The unstructured part of the HsOrc6 is between the site of attachment to the rest of ORC and these domains and is only predicted to be ∼45 aa long. This shorter length could make formation of a tether more difficult, although the presence of the full interacting region of the two TFIIB-domains of Orc6 at the N-terminus could make the interaction more robust. Mutations in either the N-terminal region or C-terminal helix of HsOrc6 that lead to Meier-Gorlin syndrome have a similar defect in helicase loading to reactions lacking ORC6 (Yang et al., 2024), suggesting that the role of Orc6 is important for proper DNA replication and cellular function, potentially through this one-ORC, MO-dependent mechanism. Regardless, it will require more direct experiments to resolve whether a one-ORC mechanism can operate and it is certain that it will not be the only mechanism that contributes to origin licensing, as reactions without Orc6 are possible.

## Materials and Methods

### Nomenclature for fluorescently-modified proteins

We use a shorthand notation to describe proteins with different modifications (for example, ORC^6N-550^). The numerical superscripts 550 and 650 correspond to the fluorescent dyes DyLight 550 and DyLight 650. The site of modification is preceding the numerical superscripts; for example, ORC^6N-550^ refers to ORC labeled at Orc6-N terminal region with DyLight 550.

### Growth of cells for protein purification

Growth of cells for protein purification was performed as described (Frigola et al., 2013; Kang et al., 2014). In brief*, S. cerevisiae* strains were grown to OD_600_ = 1.0 in 6L of YEP supplemented with 2% glycerol (w/v) at 30°C. Cells were arrested in G1 using α-factor (100 ng/mL) and expression of proteins was induced using 2% galactose (w/v). After induction, cells were harvested and sequentially washed with 100 mL wash buffer and 50 mL lysis buffer. The washed pellet was resuspended in approximately 1/3 of packed cell volume with lysis buffer containing cOmplete Protease Inhibitor Cocktail Tablet (1 tablet per 25 mL total volume; Roche) and frozen dropwise in liquid nitrogen. Frozen cells were lysed in a SamplePrep freezermill (SPEX) and the powder was saved at −80°C. Upon thawing of the cell powder, the lysate was clarified by ultracentrifugation in a Type 45Ti rotor at 36 krpm (150,000 x g) for 1 hr at 4°C. This lysate was then used for further steps of protein purification as described below.

### Preparation of unlabeled proteins

Wild-type Mcm2-7, Cdc6 and Cdt1 were purified as described previously (Frigola et al., 2013; Kang et al., 2014). Clb5-Cdk1 was purified from ySK119 and Sic1 was purified from BL21-DE3-Rosetta bacteria transformed with the plasmid pGEX-Sic1 as described previously (Heller et al., 2011).

### Preparation of labeled ORC^6N-550^, ORC^6-lnk1scr-550^

For ORC^6N-550^ and ORC^6-lnk1scr-550^, an S6 tag (GDSLSWLLRLLN) along with 3 x GGS flexible linker on either side of the tag was inserted after amino acid 107 in Orc6. Dy550-CoA dyes were synthesized as described (Yin et al., 2006) using maleimide-Dylight 550 (Thermo Scientific) and Coenzyme A trilithium salt (Research Products International). ORC was purified from lysate using FLAG affinity resin (Sigma-Aldrich) as previously described (Gupta et al., 2021). ORC, SFP synthase (NEB), and Dy550-CoA were incubated in a 1:4:10 ratio for 45 minutes at room temperature, then at 4°C overnight. Labeled ORC was then purified on a Superdex 200 Increase 10/300 gel filtration column. Peak fractions were aliquoted and stored at −80°C.

### Preparation of labeled ORC^6ΔN-550^, ORC^6ΔN-lnk1scr-550^, ORC^6C-550^

For sortase-mediated labeling, peptides (NH2-CHHHHHHHHHLPETGG-COOH for N-terminal lableing, NH2-GGGHHHHHHHHHHC-COOH for C-terminal labeling) were labeled with maleimide-derivatized Dylight 550 or Dylight 650 and HPLC purified.

ORC was purified from lysate using FLAG affinity resin (Sigma-Aldrich) as previously described (Gupta et al., 2021). Sortase-mediated labeling was performed by incubating purified protein with equimolar amount of Sortase along with 100 nmol of either the labeled N-or C-peptide in buffer supplemented with 5 mM CaCl_2_ for 15 minutes at room temperature. After incubation, the reaction was quenched with 15 mM EDTA and applied to a Superdex 200 Increase 10/300 gel filtration column to remove Sortase and unreacted peptide-dye. Peak protein fractions were incubated with 0.3 mL of Ni-NTA Agarose Resin (Qiagen) in buffer supplemented with 5 mM imidazole overnight at 4°C. The resin was collected and then washed first with 5 mL buffer supplemented with 15 mM imidazole, then washed with 5mL buffer supplemented with 25 mM imidazole. Labeled protein was then eluted in buffer supplemented with 300 mM imidazole. Peak fractions were aliquoted and stored at −80°C.

### Preparation of labeled Mcm2-7^6N-650^ and Mcm2-7^2C-650^

Mcm2-7 protein was purified using FLAG resin and labeled via Sortase-mediated conjugation as described above. For Mcm2-7^6N-650^, the N-terminal extension of Mcm6 (2-103) (which is not important for helicase loading, Champasa et al. 2019) was deleted and replaced with an N-terminal Sortase recognition sequence Ubiquitin-GGG. During expression in cells, the ubiquitin is cleaved from the protein, and after Sortase labeling the N-terminal Sortase peptide coupled to Dylight-650 is attached to the N-terminus of Mcm6Δ103. For Mcm2-7^2C-650^, a C-terminal Sortase recognition sequence (LPETGG) was placed at the C-terminus of Mcm2. During Sortase labeling, the C-terminal Sortase peptide coupled to Dylight-650 is attached to the C-terminus of Mcm2.

### Preparation of labeled Mcm2-7^3N-650^ and Cdt1^C-650^

Mcm2-7^3N-650^ and Cdt1^C-650^ were purified as described (Gupta et al., 2021; Ticau et al., 2015).

### Single-molecule assay for helicase loading

A micro-mirror total internal reflection microscope was used to perform multiwavelength single-molecule imaging (Friedman et al., 2006). Glass slides were functionalized with Biotin-PEG and PEG, and biotinylated DNA along with streptavidin-coated fiducial markers (0.04 micron, ThermoFisher TransFluoSpheres T10711) were coupled to the slide as described (Gupta et al., 2021). All reactions were performed as described previously (Kang et al., 2014; Ticau et al., 2015) in buffer containing 25 mM HEPES-KOH pH 7.6, 300 mM potassium glutamate, 5 mM Mg(OAc)_2_, 3 mM ATP, 1mM dithiothreitol, 1 mg/mL bovine serum albumin, with an oxygen scavenging system (glucose oxidase/catalase) as well as 2 mM Trolox (Crawford et al., 2008). Reactions used 0.5-1 nM ORC, 3-10 nM Cdc6, and 10-15 nM Mcm2-7/Cdt1. All reactions also included 0.5 µM of a 60 bp nonspecific DNA generated by annealing the following two oligonucleotides: (F: 5-CTTGTTATTTTACAGATTTTCTCCATTCTTCTTTTATGCTTGCAAAACAAAAGGCCTGCA-3’ R:5’-TGCAGGCCTTTTGTTTTGCAAGCATAAAAGAAGAATGGAGAAAATCTGTAAAATAACAAG-3). Positions of DNA molecules labeled with Alexa Fluor 488 were located before experiments using 488 nM excitation. Experimental acquisition was performed using 1 s alternating 532 nm and 633 nm excitation, with approximately 0.2 s of dead time between acquisitions.

### FRET Data Analysis

CoSMoS data sets were analyzed as described previously (Ticau et al., 2015) and fluorescence intensity values were background corrected as described (De Jesús-Kim et al., 2021). Apparent *E*_FRET_ calculations performed as described (Gupta et al., 2021). To determine identity of low and high *E*_FRET_ states, the *E*_FRET_ data for ORC and Mcm2-7^6N-650^ interactions (experiments in Figure S2) were selected from two time intervals after first Mcm2-7^6N-650^ arrival for each experiment: 0 to 5 s and >5 s. *E*_FRET_ values from −0.2 to 1.2 (which constitute >97% of observations) from each interval were independently fit to the two-component Gaussian mixture probability density function:

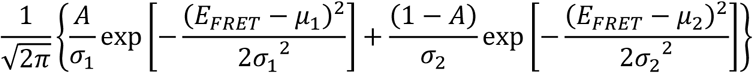

where A is the fractional amplitude of the low *E*_FRET_ component, *μ_1_* < *μ*_2_ are the mean *E*_FRET_ values of the low and high components, and *σ_1_*, *σ_2_* are the SDs of the low and high *E*_FRET_ components. SEs of the fit parameters were computed by bootstrapping (1,000 samples). The *E*_FRET_ threshold value used to differentiate between the low and high *E*_FRET_ state was defined as the crossing point of the two components of the gaussian fit for each dataset. An *E*_FRET_ transition from low to high was considered to have taken place once the *E*_FRET_ value crossed the threshold for two consecutive frames. An *E*_FRET_ transition from high to low was considered to have taken place once the *E*_FRET_ value crossed below the threshold for two consecutive frames. Median time to first formation of interaction (Figure S1E, 2C, 4C, S7C) was determined by taking the median time at which the high *E*_FRET_ state was reached for all molecules that successfully made a transition from low to high *E*_FRET_. Median duration of interaction (Figure S1F, 2D, 4D, S7F) was determined by taking the median time of the duration of the high *E*_FRET_ states for all molecules that made a transition from low to high *E*_FRET_ during the experiment.

### *E*_FRET_ heat maps

*E*_FRET_ heat maps were generated as described (Zhang et al., 2023). In brief, heat maps were constructed using MATLAB code adapted from https://github.com/gelles-brandeis/jganalyze. The code uses two-dimensional kernel density estimation, with a normal kernel function (standard deviation of time axis = 5s and *E*_FRET_ axis 0.05) to generate a heat map of *E*_FRET_ values versus time (0-40 seconds after first Mcm2-7 arrival). Each vertical slice represents the distribution of *E*_FRET_ values at that particular time interval. The time axis has a resolution of 1 second and the *E*_FRET_ axis has a resolution of 0.005. Normalization was performed such that the density estimate at each time slice integrates to 1.

### Ensemble helicase loading assays

Ensemble helicase loading assays (Figure S1B, S8) were performed as described previously (Kang et al., 2014).

### Plasmid shuffle assays

Plasmid shuffle assays to test viability of Orc6 mutants (Figure 2F) were performed essentially as described previously (Sikorski and Boeke, 1991). Briefly, a yeast strain (ySC136) with the chromosomal copy of *ORC6* deleted and a plasmid containing a WT copy of *ORC6* and a *URA3* marker were transformed with an integrating plasmid expressing the indicated test allele of *ORC6*. Transformed cells were grown in YPD liquid media overnight at 30°C (allowing loss of the *URA3* plasmid) and plated onto either YPD or 5-FOA plates in 10-fold serial dilutions, starting with 5 µL of 0.5 OD_600_ cells in the first spot. Plates were incubated at 30°C for 2 days before imaging.

## Acknowledgments

This work was supported by NIH grants R01 GM147960 (S.P.B. and J.G) and R01 GM81648 (J.G). S.P.B. is an investigator with the Howard Hughes Medical Institute. This work was supported in part by the Koch Institute Support Grant P30-CA14051 from the NCI. We thank the Koch Institute Swanson Biotechnology Center for technical support. We thank Brett Kahmann for help with protein production.

