## Supplemental Figures and Tables for "An Orc6 tether mediates ORC binding site switching during replication origin licensing"

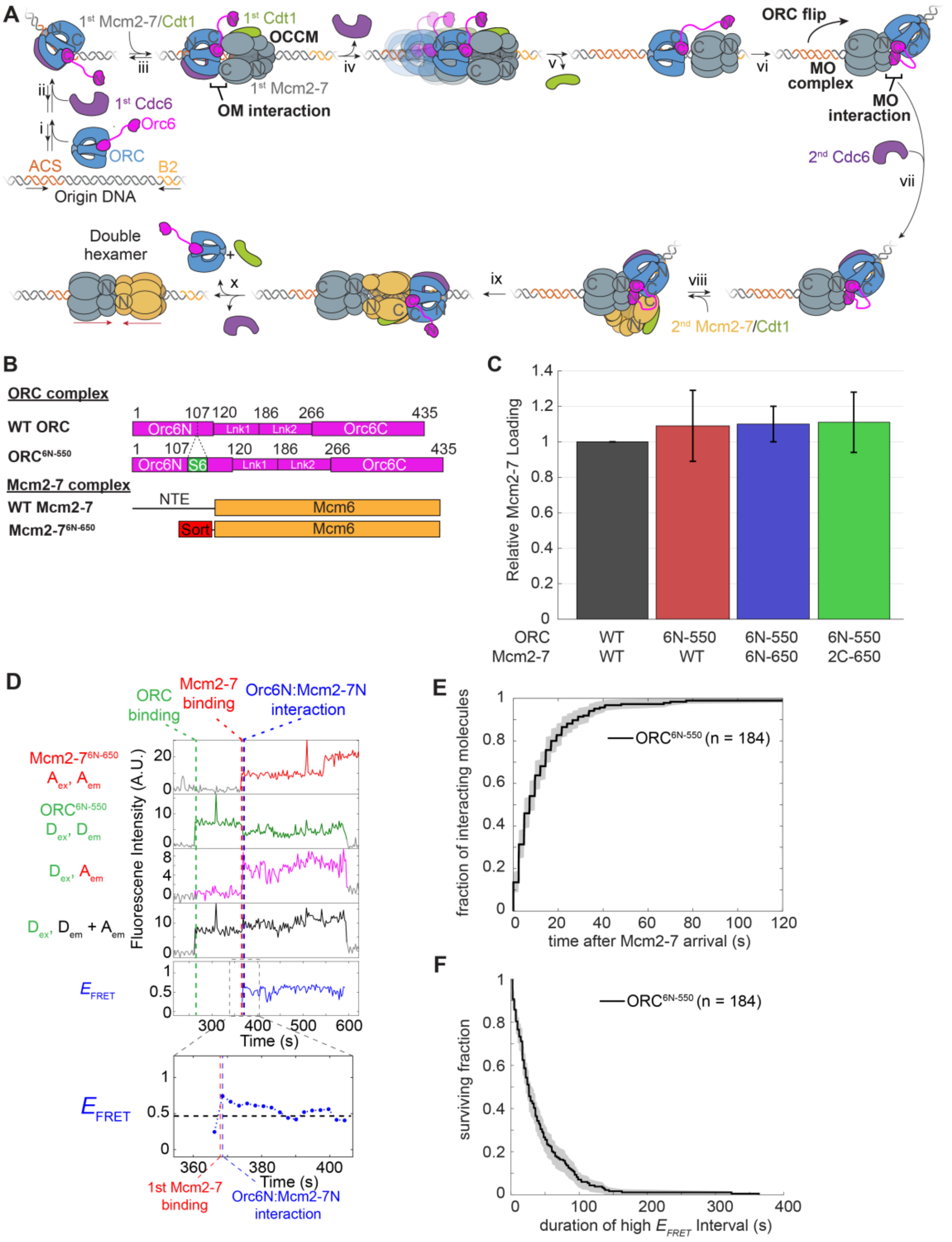


**Figure S1**: Helicase loading model prior to this study and additional information about assay.

A: Model of Helicase Loading prior to data from this study. See text for description. N and C notations on ORC and Mcm2-7 represent the N- and C-tiers of each protein. N and C notations on Orc6 represent the N- and C-terminal TFIIB-related domains.

B: Labeling approach for ORC$^{6N-550}$ and Mcm2-7$^{6N-650}$. ORC$^{6N-550}$ was modified by inserting an S6 peptide tag (GDSLSWLLRLLN) after amino acid 107 in Orc6. ORC containing the modified Orc6 was purified and labeled using Sfp synthase and acetyl-CoA Dylight 550. For Mcm2-7$^{6N-650}$, Mcm6 was modified by replacing amino acids 1-103 (NTE) with an N-terminal recognition tag for Sortase. Mcm2-7 containing the modified Mcm6 was labeled using Sortase to attach Dylight 650-maleimide conjugated peptide.

C: Ensemble helicase-loading assays were used to assess ORC$^{6N-550}$, Mcm2-7$^{6N-650}$, and Mcm2-7$^{2C-650}$ function. SDS-gel band intensities were quantified relative to parallel reactions with unmodified wild-type ORC and Mcm2-7 to obtain relative extents of Mcm2-7 loading. Errors are SEM. N = 2 for each reaction.

D: Additional record showing Mcm2-7$^{6N-650}$ recruitment to ORC$^{6N-550}$ resulting in double hexamer formation plotted as in Figure 1C. This record illustrates that the high $E_{FRET}$ state does not have to be maintained at all times for successful double-hexamer formation.

E: Time to formation of the Orc6N:Mcm2-7N interaction relative to Mcm2-7$^{6N-650}$ arrival. Horizontal axis is time after Mcm2-7$^{6N-650}$ recruitment. Vertical axis is the fraction of stable ORC-Mcm2-7 complexes that formed the Orc6N:Mcm2-7N interaction. Shading represents 95% CI. Only the first Orc6N:Mcm2-7N interaction for a given recruitment of Mcm2-7$^{6N-650}$ by ORC$^{6N-550}$ was considered for this analysis. Molecules that did not achieve a high $E_{FRET}$ state were not included in the analysis.

F: Cumulative survival curve of Orc6N:Mcm2-7N interaction. This plot shows the distribution of lifetimes for the Orc6N-Mcm2-7N interactions. The same subset of events included in E are presented in this analysis. Shading represents 95% CI.

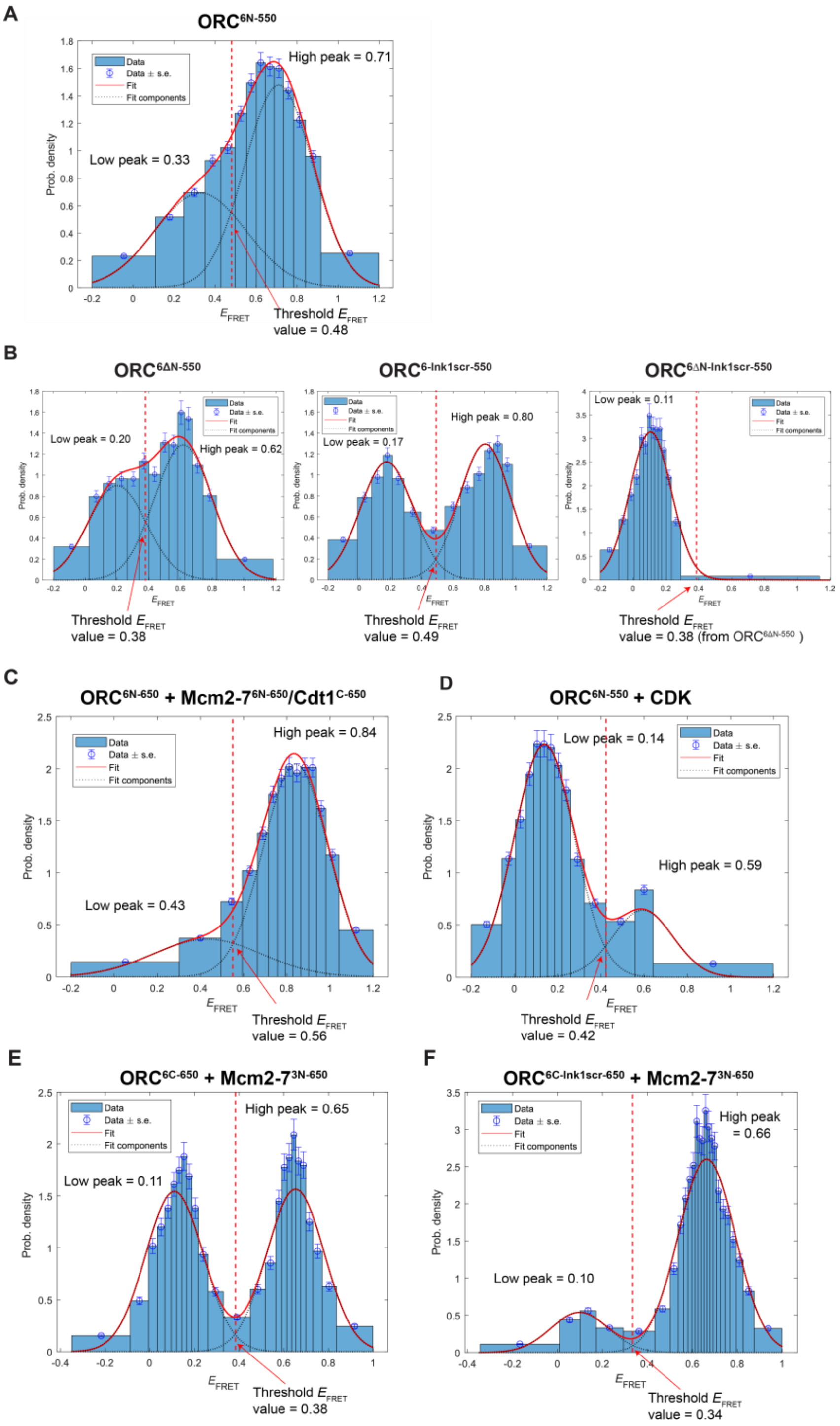


**Figure S2**: Fits of $E_{FRET}$ distributions to two-component Gaussian mixture models (see Methods). The thresholds to delineate between low and high $E_{FRET}$ states were taken to be the points (arrows) at which the component curves intersect. Fit parameters and N values are presented in Table S1.

A: Experiment (Figure 1) with ORC$^{6N\text{-}550}$ and Mcm2-7$^{6N\text{-}650}$.

B: Experiments (Figure 2) with ORC$^{6\Delta N-550}$ and Mcm2-7$^{6N-650}$ (left), ORC$^{6-lnk1scr-550}$ and Mcm2-7$^{6N-650}$ (middle), and ORC$^{6\Delta N-lnk1scr-550}$ and Mcm2-7$^{6N-650}$ (right).

C: Experiment (Figure 3) with ORC$^{6N-550}$ and Mcm2-7$^{6N-650}$ / Cdt1$^{C-650}$.

D: Experiment (Figure 4) with ORC$^{6N-550}$, Mcm2-7$^{6N-650}$, and CDK.

E: Experiment (Figure 5) with ORC$^{6C-550}$ and Mcm2-7$^{3N-650}$.

F: Experiment (Figure 5) with ORC$^{6C-lnk1scr-550}$ and Mcm2-7$^{3N-650}$.

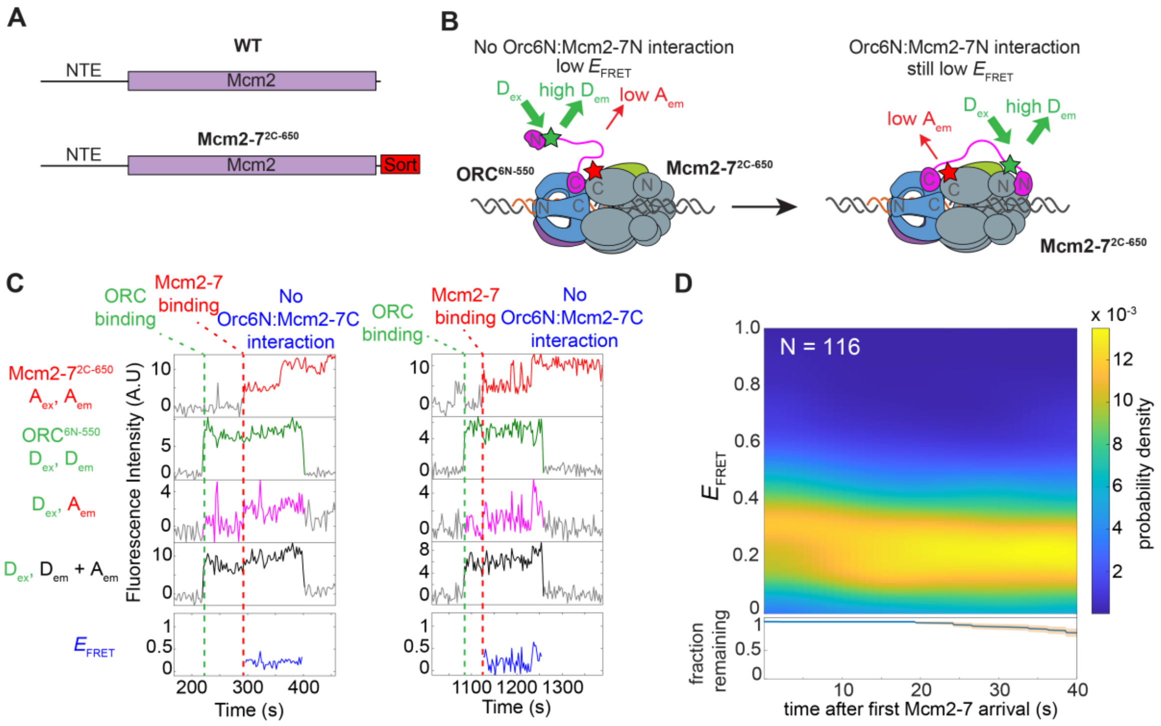


**Figure S3**: ORC$^{6N-550}$ FRET with Mcm2-7 is specific to the Mcm2-7 N-terminus.

A: Labeling approach for Mcm2-7$^{2C-650}$. Mcm2 was modified by attaching a C-terminal recognition tag for Sortase. Mcm2-7 complexes with the Sortase tags were coupled to a Dylight-650-labeled peptide using Sortase. See Methods for details.

B: Model of FRET observed in an experiment in which ORC$^{6N-550}$ was labeled with donor fluorophore as in Figure 1 but Mcm2-7$^{2C-650}$ was labeled with an acceptor fluorophore at the C-terminus of Mcm2.

C: Two representative single-DNA records of ORC$^{6N-550}$ recruitment of Mcm2-7$^{2C-650}$, plotted as in Figure 1C.

D: $E_{FRET}$ distribution heat map for 116 DNA molecules where ORC$^{6N-550}$ recruited Mcm2-7$^{2C-650}$. Bottom plot shows fraction of ORC$^{6N-550}$-Mcm2-7$^{2C-650}$ complexes that retain both ORC$^{6N-550}$ and Mcm2-7$^{2C-650}$ bound and the 95% CI are shown in the bottom plot (blue curve, orange shading).

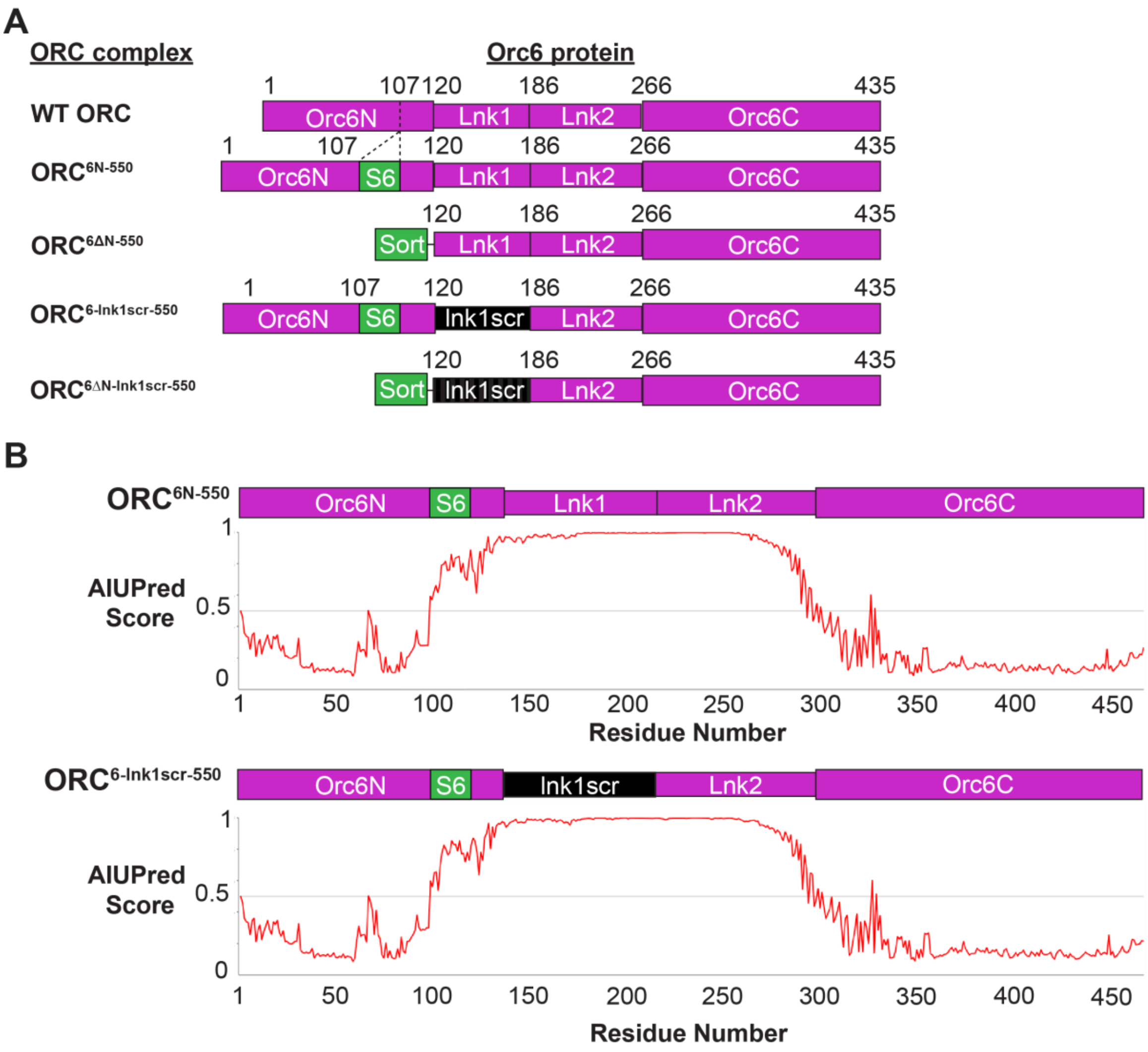


**Figure S4**: ORC constructs used for Orc6 mutant experiments.

A: Diagram of Orc6 mutants used in Orc6-tether assays. For each ORC complex tested in this study, mutations and the modifications made for labeling of Orc6 protein are shown. $ORC^{6N\text{-}550}$ and $ORC^{6\text{-}lnk1scr\text{-}550}$ were labeled with the Sfp synthase/S6 tag system described in Figure S1B. $ORC^{6\Delta N\text{-}550}$ and $ORC^{6\Delta N\text{-}lnk1scr\text{-}550}$ were labeled using Sortase as described in Figure S1B. See Methods for protein labeling details.

B: Structure prediction of amino acids in $ORC^{6N\text{-}550}$ and $ORC^{6\text{-}lnk1scr\text{-}550}$ using AIUpred (Erdős and Dosztányi, 2024). Values greater than 0.5 indicate regions predicted to be disordered.

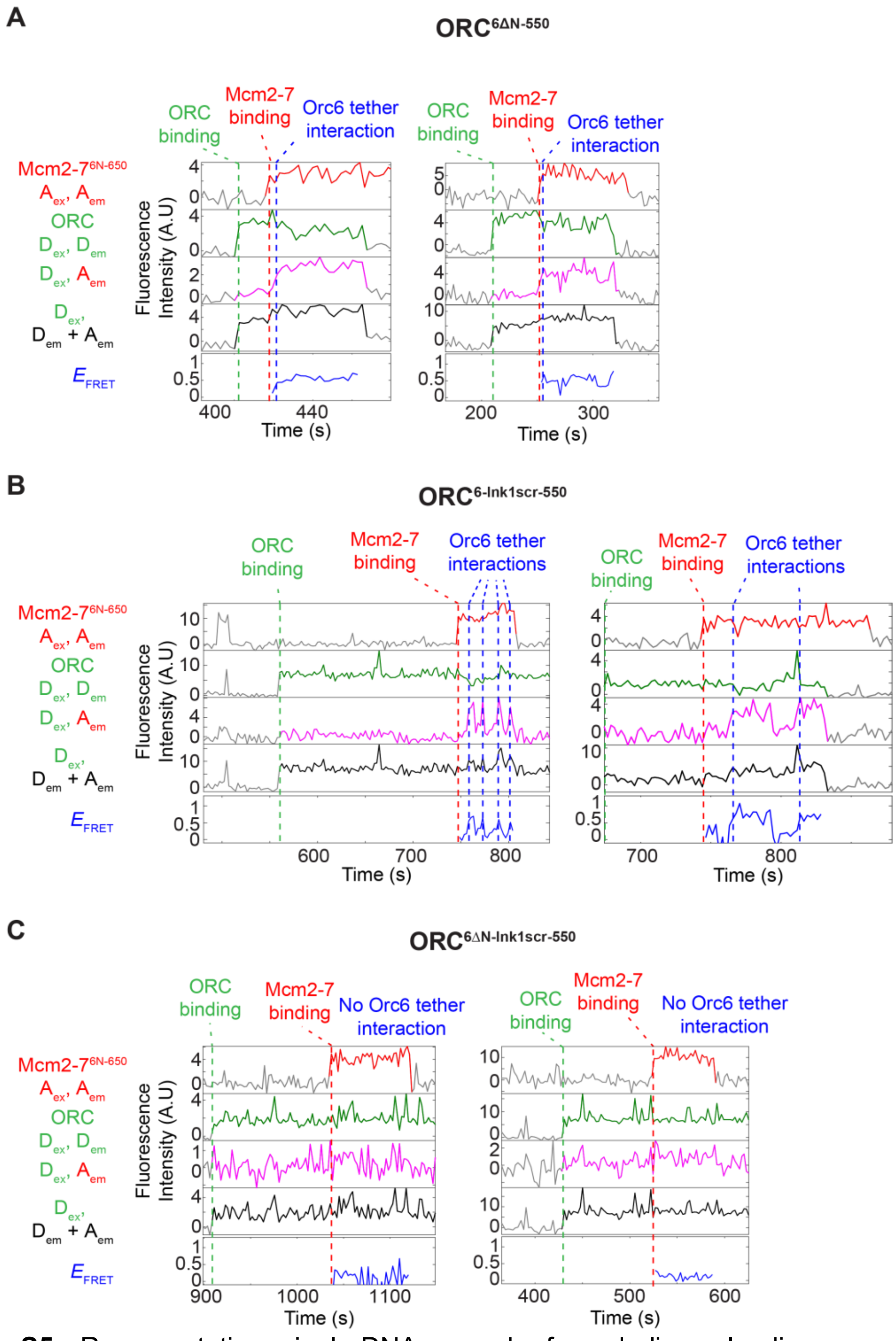


**Figure S5**: Representative single-DNA records from helicase-loading experiments described in Figure 2 using Mcm2-$7^{6N\text{-}650}$ and $ORC^{6\Delta N\text{-}550}$ (A), $ORC^{6\text{-}Ink1scr\text{-}550}$ (B), or $ORC^{6\Delta NInk1scr550}$. Records are plotted as in Figure 1C.

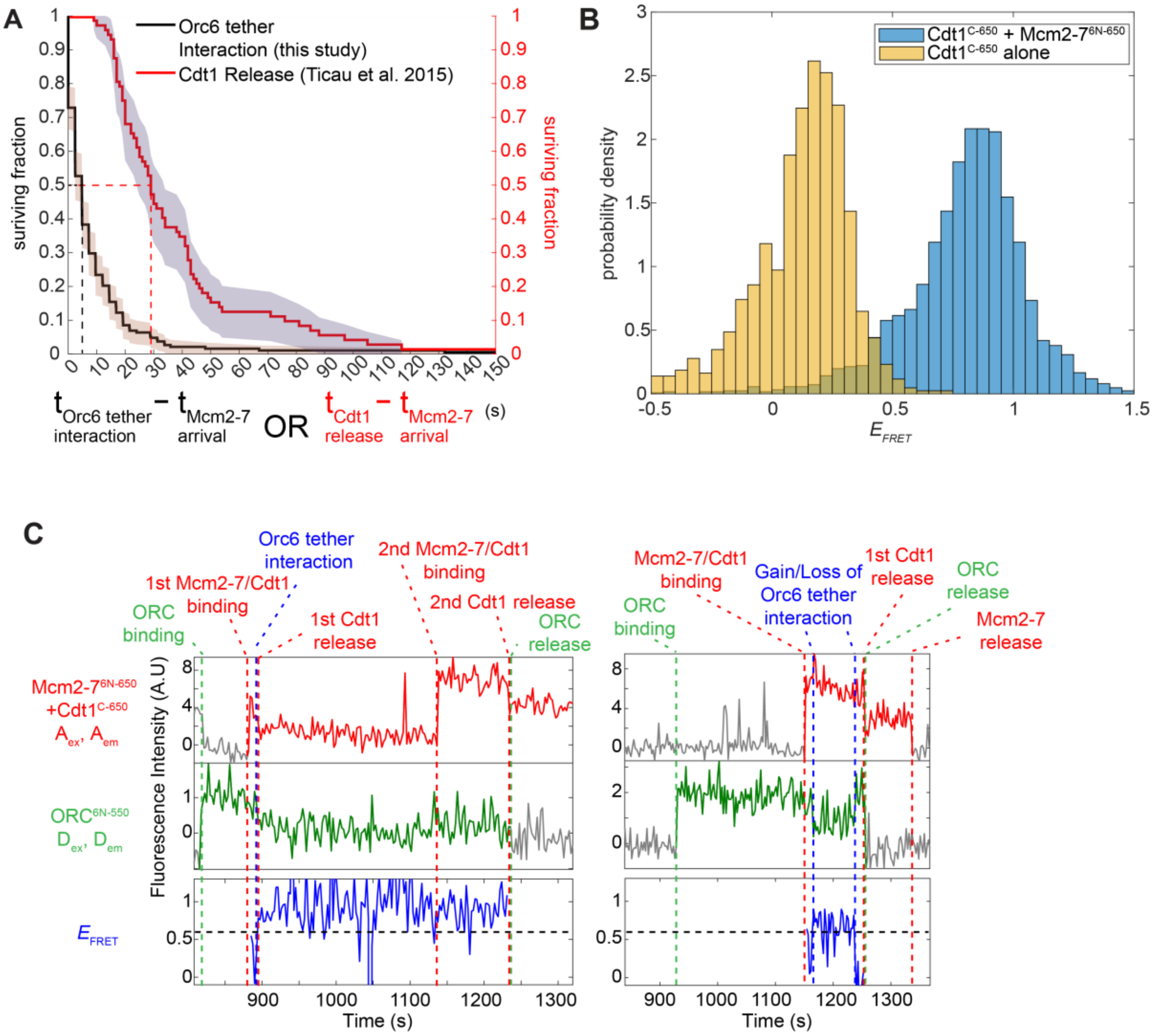


**Figure S6**: Timing of Orc6 tether interaction and Cdt1 release.

A: Survival plot of the fraction of molecules with Cdt1 (red) or that have yet to form the Orc6 tether interaction (black). All times are relative to arrival of first Mcm2-7 at the same DNA molecule. Shaded areas represent 95% CI. Dashed lines indicate median times for each dataset.

B: $Cdt1^{C\text{-}650}$ does not exhibit high $E_{\mathrm{FRET}}$ with $ORC^{6N\text{-}550}$ during helicase loading. Histogram plots of $E_{\mathrm{FRET}}$ values from $Mcm2\text{-}7^{6N\text{-}650}$ recruitment events during each frame of colocalization of ORC and Mcm2-7/Cdt1 to DNA. Yellow bars: $Cdt1^{C\text{-}650}$, $ORC^{6N\text{-}550}$ and unlabeled Mcm2-7 and Cdc6. Blue bars: $Cdt1^{C\text{-}650}$, $ORC^{6N\text{-}550}$, $Mcm2\text{-}7^{6N\text{-}650}$ and unlabeled Cdc6.

C: Additional traces of Orc6 tether interaction and Cdt1-association assay using $ORC^{6N\text{-}550}$, $Mcm2\text{-}7^{6N\text{-}650}$ and $Cdt1^{C\text{-}650}$. The right panel shows an example in which $E_{\mathrm{FRET}}$ is low at the time of 1st Cdt1 release and, consistent with a tether function, ORC release occurs rapidly after 1st Cdt1 release. Traces are arranged as described in Figure 3C.

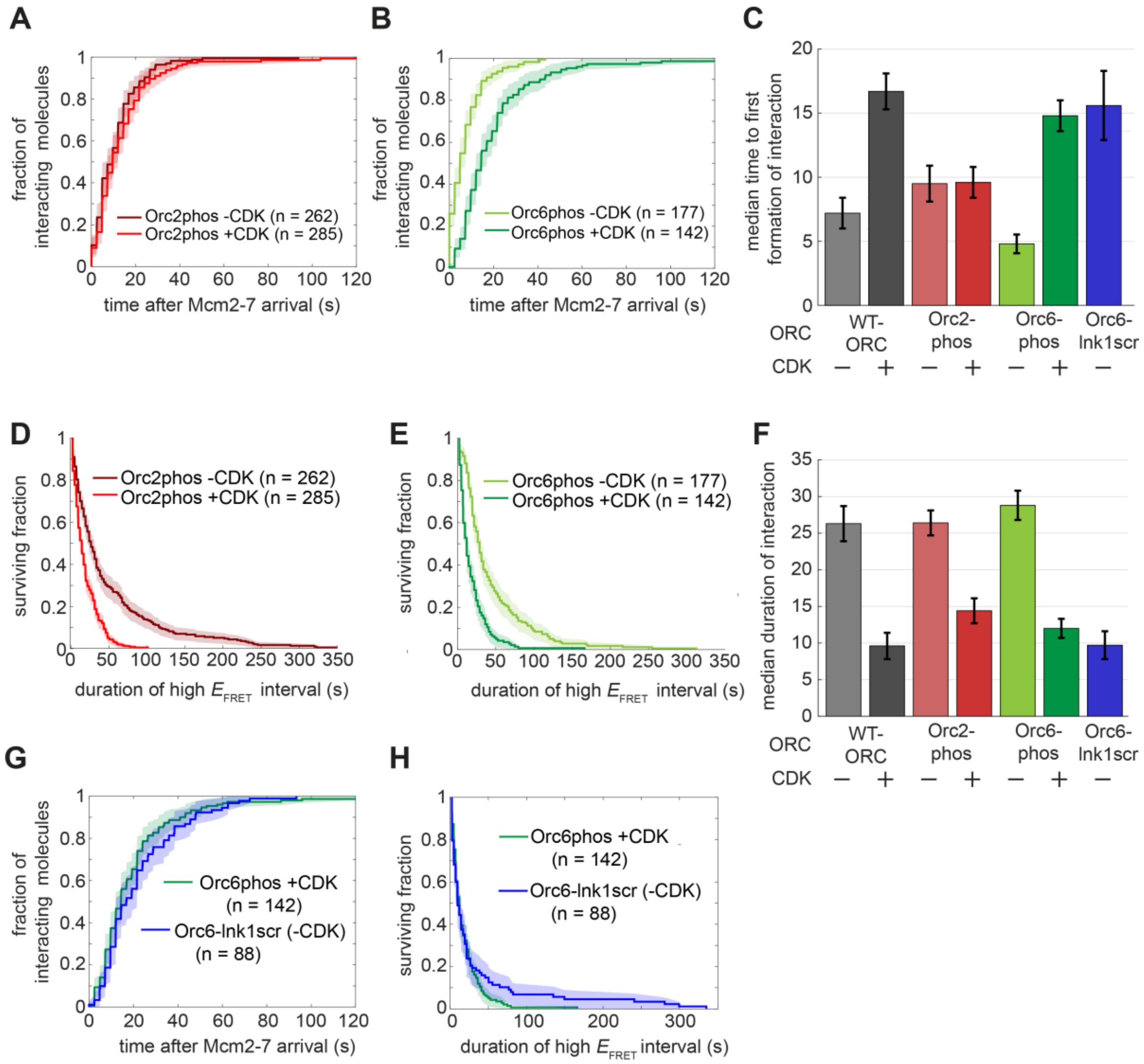


**Figure S7**: Phosphorylation of Orc6 alone inhibits median time to formation of Orc6 tether, but phosphorylation of either Orc2 or Orc6 impacts the stability of the Orc6 tether interaction.

A: Plot same as Figure 4C but using ORC$^{2phos\text{-}6N\text{-}550}$.

B: Plot same as Figure 4C but using ORC$^{6phos\text{-}6N\text{-}550}$.

C: Median time to first tether interaction formation with and without CDK modification for WT, ORC$^{2phos\text{-}6N\text{-}550}$, and ORC$^{6phos\text{-}6N\text{-}550}$. The time to first formation (± S.E.) for unmodified ORC$^{6\text{-}Ink1scr\text{-}650}$ (Figure 2C) is shown for comparison.

D: Plot same as Figure 4D but using ORC$^{2phos\text{-}6N\text{-}550}$.

E: Plot same as Figure 4D but using ORC$^{6phos\text{-}6N\text{-}550}$.

F: Median duration of first tether interaction with and without CDK modification for WT, $\text{ORC}^{\text{2phos-6N-550}}$, and $\text{ORC}^{\text{6phos-6N-550}}$. Median duration of first tether interaction (± S.E.) for unmodified $\text{ORC}^{\text{6-Ink1scr-650}}$ (Figure 2D) is shown for comparison.

G: Plot same as Figure 4C but now comparing $\text{ORC}^{\text{6phos-6N-550}}$ +CDK (green, same data as B) to $\text{ORC}^{\text{6-Ink1scr-550}}$ (blue, same data as Figure 2C).

H: Cumulative survival curve of initial Orc6 tether interactions for CDK-modified $\text{ORC}^{\text{6phos-6N-550}}$ (green) or unmodified $\text{ORC}^{\text{6-Ink1scr-550}}$ (blue). Shading represents 95% CI. Note, ~20% of $\text{ORC}^{\text{6-Ink1scr-550}}$ molecules retain the tether interaction for significantly longer times compared to $\text{ORC}^{\text{6phos-6N-550}}$ +CDK.

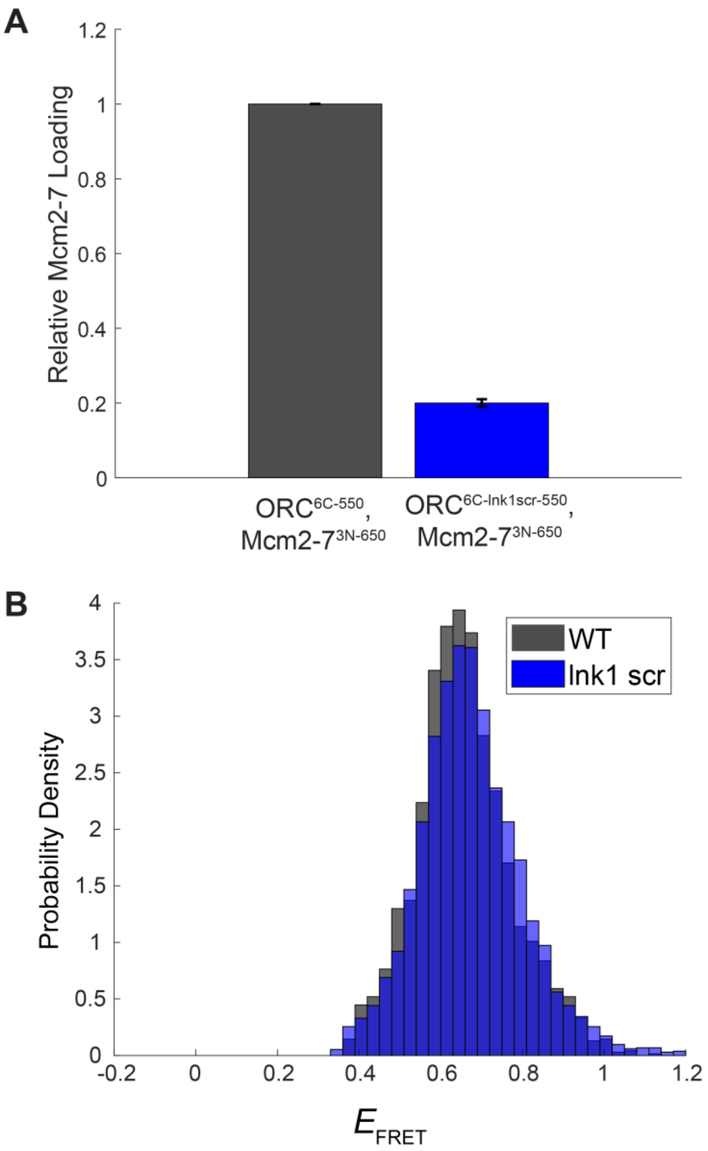


**Figure S8**: Further analysis of Orc6 Ink1 function.

A: $ORC^{6C\text{-}Ink1scr\text{-}550}$ is deficient in double-hexamer formation in ensemble helicase-loading assays. Ensemble helicase-loading assays were performed with $ORC^{6C\text{-}550}$ or $ORC^{6C\text{-}Ink1scr\text{-}550}$ and $Mcm2\text{-}7^{3N\text{-}650}$, along with unlabeled Cdt1 and Cdc6. Bars indicate the relative loading of salt-stable helicases. Errors are S.E.M. N = 2 for each reaction.

B: $E_{FRET}$ values of MO complex for $ORC^{6C\text{-}550}$ and $ORC^{6C\text{-}Ink1scr\text{-}550}$ are similar. $E_{FRET}$ values from each experiment (black: $ORC^{6C\text{-}550}$, blue: $ORC^{6C\text{-}Ink1scr\text{-}550}$) were computed and binned into low and high $E_{FRET}$ states based on threshold determination (see Figure S2E-F, Methods). The resulting high $E_{FRET}$ values were plotted as a histogram and normalized by probability density to compare the distribution of high $E_{FRET}$ values.

**Table S1. Fit parameters for $E_{FRET}$ distributions***

| ORC | Mcm2-7 | Cdt1 | Figure | A | μ1 | σ1 | μ2 | σ2 | N |
|---|---|---|---|---|---|---|---|---|---|
| $ORC^{6N\text{-}550}$ | $Mcm2\text{-}7^{6N\text{-}650}$ | WT | 1 | 0.384 | 0.33 | 0.22 | 0.71 | 0.17 | 7613 |
| $ORC^{6\Delta N\text{-}550}$ | $Mcm2\text{-}7^{6N\text{-}650}$ | WT | 2 | 0.414 | 0.20 | 0.18 | 0.62 | 0.18 | 3226 |
| $ORC^{6\text{-}lnk1scr\text{-}550}$ | $Mcm2\text{-}7^{6N\text{-}650}$ | WT | 2 | 0.473 | 0.17 | 0.17 | 0.80 | 0.16 | 4239 |
| $ORC^{6\Delta N\text{-}lnk1scr\text{-}550}$ | $Mcm2\text{-}7^{6N\text{-}650}$ | WT | 2 | 0.991 | 0.11 | 0.12 | 0.57 | 0.30 | 3137 |
| $ORC^{6N\text{-}550}$ | $Mcm2\text{-}7^{6N\text{-}650}$ | $Cdt1^{C\text{-}650}$ | 3 | 0.228 | 0.43 | 0.25 | 0.84 | 0.15 | 7680 |
| $ORC^{6N\text{-}550}$ + CDK | $Mcm2\text{-}7^{6N\text{-}650}$ | WT | 4 | 0.776 | 0.14 | 0.14 | 0.59 | 0.14 | 4894 |
| $ORC^{6C\text{-}550}$ | $Mcm2\text{-}7^{3N\text{-}650}$ | WT | 5 | 0.503 | 0.11 | 0.13 | 0.65 | 0.13 | 5490 |
| $ORC^{6C\text{-}lnk1scr\text{-}550}$ | $Mcm2\text{-}7^{3N\text{-}650}$ | WT | 5 | 0.868 | 0.10 | 0.13 | 0.66 | 0.13 | 6089 |

*A, proportion of data in low $E_{FRET}$ component; μ1, mean of low $E_{FRET}$ component; σ1, SD of low $E_{FRET}$ component; μ2, mean of high $E_{FRET}$ component; σ2, SD of high $E_{FRET}$ component; N = number of observations for fits.

**Table S2. List of yeast strains used in this study**

| Strain Name | Construct | Genotype | Study |
|---|---|---|---|
| yDD57 | $ORC^{6N-550}$ | *ade2-1 trp1-1 leu2-3,112 his3-11,15 ura3-1 can1-100 bar1::hisG lys2::HisG pep4::unmarked URA3::pAZ96 (GAL1,10 FLAG-ORC1-opt, ORC2-opt) HIS3::pJF17 (ORC3-opt, ORC4-opt) TRP1::pDD48 (GAL1,10 ORC5, ORC6-107-S6)* | This study |
| yDD50 | Mcm2-$7^{6N-650}$/Cdt1 | *ade2-1 trp1-1 leu2-3,112 his3-11,15 ura3-1 can1-100 bar1::hisG lys2::HisG pep4::unmarked LYS2::pSKM002 (GAL1,10 MCM4, MCM5) TRP1::pDD36 (GAL1,10 UbSORT-mcm6∆103, MCM7) HIS3::pSKM004 (GAL1,10 MCM2, Flag-MCM3) URA3::pALS1 (GAL1,10 CDT1, GAL4)* | This study |
| ySG01 | Mcm2-$7^{2C-650}$/Cdt1 | *ade2-1 trp1-1 leu2-3,112 his3-11,15 ura3-1 can1-100 bar1::hisG lys2::HisG pep4::unmarked LYS2::pSKM002 (GAL1,10 MCM4, MCM5) TRP1::pSKM003 (GAL1,10 MCM6, MCM7) HIS3::pST048 (GAL1,10 MCM2C-LPETGG, Flag-MCM3) URA3::pALS1 (GAL1,10 CDT1, GAL4)* | Gupta et al. 2021 |
| yAZ90 | $ORC^{6\Delta N-550}$ | *ade2-1 trp1-1 leu2-3,112 his3-11,15 ura3-1 can1-100 bar1::hisG lys2::HisG pep4::unmarked URA3::pAZ96 (GAL1,10 FLAG-ORC1-opt, ORC2-opt) HIS3::pJF17 (ORC3-opt, ORC4-opt) TRP1::pAZ87 (GAL1,10 ORC5, UbSORT-orc6ΔN)* | This study |
| yDD54 | $ORC^{6-lnk1scr-550}$ | *ade2-1 trp1-1 leu2-3,112 his3-11,15 ura3-1 can1-100 bar1::hisG lys2::HisG pep4::unmarked URA3::pAZ96 (GAL1,10 FLAG-ORC1-opt, ORC2-opt) HIS3::pJF17 (ORC3-opt, ORC4-opt) TRP1::pDD53 (GAL1,10 ORC5, orc6-107-S6-lnk1scr)* | This study |

| | | | |
|---|---|---|---|
| yDD52 | ORC$^{6\Delta N\text{-lnk1scr-550}}$ | *ade2-1 trp1-1 leu2-3,112 his3-11,15 ura3-1 can1-100 bar1::hisG lys2::HisG pep4::unmarked URA3::pAZ96 (GAL1,10 FLAG-ORC1-opt, ORC2-opt) HIS3::pJF17 (ORC3-opt, ORC4-opt) TRP1::pDD49 (GAL1,10 ORC5, UbSORT-orc6ΔN-lnk1scr)* | This study |
| ySC136 | ORC6 swapper strain | *ade2-1 ura3-1 his3-11,15 trp1-1 leu2-3,112 can1-100 lys2::hisG bar1::hisG orc6::KanMX MATa pSPB66 (ORC6, URA3)* | Chen et al. 2011 |
| ySC154 | Orc3pro-ORC6 | *ade2-1 ura3-1 his3-11,15 trp1-1 leu2-3,112 can1-100 lys2::hisG bar1::hisG orc6::KanMX MATa pSPB66 (ORC6, URA3) TRP1::pRS404-Orc3p-ORC6* | Chen et al. 2011 |
| yDD70 | Orc3pro-*orc6ΔN* | *ade2-1 ura3-1 his3-11,15 trp1-1 leu2-3,112 can1-100 lys2::hisG bar1::hisG orc6::KanMX MATa pSPB66 (ORC6, URA3) TRP1::pDD73 (Orc3p-orc6ΔN)* | This study |
| yDD71 | Orc3pro-*orc6-lnk1scr* | *ade2-1 ura3-1 his3-11,15 trp1-1 leu2-3,112 can1-100 lys2::hisG bar1::hisG orc6::KanMX MATa pSPB66 (ORC6, URA3) TRP1::pDD74 (Orc3p-orc6-lnk1scr)* | This study |
| yDD72 | Orc3pro-*orc6ΔN-lnk1scr* | *ade2-1 ura3-1 his3-11,15 trp1-1 leu2-3,112 can1-100 lys2::hisG bar1::hisG orc6::KanMX MATa pSPB66 (ORC6, URA3) TRP1::pDD75 (Orc3p-orc6ΔN-lnk1scr)* | This study |
| yDD46 | Mcm2-7$^{6N\text{-}650}$ (no Cdt1) | *ade2-1 trp1-1 leu2-3,112 his3-11,15 ura3-1 can1-100 bar1::hisG lys2::HisG pep4::unmarked LYS2::pSKM002 (GAL1,10 MCM4, MCM5) TRP1::pDD36 (GAL1,10 UbSORT-mcm6Δ103, MCM7) HIS3::pSKM004 (GAL1,10 MCM2, Flag-MCM3)* | This study |
| ySG46 | Cdt1$^{C\text{-}650}$ | *ade2-1 trp1-1 leu2-3,112 his3-11,15 ura3-1 can1-100 bar1::hisG lys2::HisG pep4::unmarked URA3::pSG26(GAL1,10-3xFLAG-Cdt1-C-LPETGG)* | Gupta et al 2021 |

| | | | |
|---|---|---|---|
| ySK119 | CDK | *ade2-1 trp1-1 leu2-3,112 his3-11,15 ura3-1 can1-100 bar1::HisG lys2::HisG pep4::unmarked URA3::GAL1,10 Δ2-95-CLB5-Flag CDC28-His* | Heller et al. 2011 |
| yDD73 | Orc2phos | *ade2-1 trp1-1 leu2-3,112 his3-11,15 ura3-1 can1-100 bar1::hisG lys2::HisG pep4::unmarked URA3::pAZ96 (GAL1,10 FLAG-ORC1-opt, ORC2) HIS3::pJF17 (ORC3-opt, ORC4-opt) TRP1::pDD64 (GAL1,10 ORC5, orc6-107-S6-4A)* | This study |
| yDD74 | Orc6phos | *ade2-1 trp1-1 leu2-3,112 his3-11,15 ura3-1 can1-100 bar1::hisG lys2::HisG pep4::unmarked URA3::pSG50 (GAL1,10 FLAG-ORC1-opt, orc2-6A) HIS3::pJF17 (ORC3-opt, ORC4-opt) TRP1::pDD48 (GAL1,10 ORC5, ORC6-107-S6)* | This study |
| ySG39 | $ORC^{6C\text{-}550}$ | *ade2-1 trp1-1 leu2-3,112 his3-11,15 ura3-1 can1-100 bar1::hisG lys2::HisG pep4::unmarked URA3::pJF19 (GAL1,10 CBP-ORC1-opt, ORC2-opt) HIS3::pJF17 (ORC3-opt, ORC4-opt) TRP1::pAZ63 (GAL1,10 ORC5-opt, ORC6-C-LPETGG)* | Gupta et al. 2021 |
| ySG24 | $Mcm2\text{-}7^{3N\text{-}650}$ | *ade2-1 trp1-1 leu2-3,112 his3-11,15 ura3-1 can1-100 bar1::hisG lys2::HisG pep4::unmarked LYS2::pSKM002 (GAL1,10 MCM4, MCM5) TRP1::pSKM003 (GAL1,10 MCM6, MCM7) HIS3::pSG13 (GAL1,10 MCM2, Flag-TEV-GG-MCM3)* | Gupta et al. 2021 |
| ySG54 | $ORC^{6C\text{-}lnk1scr\text{-}550}$ | *ade2-1 trp1-1 leu2-3,112 his3-11,15 ura3-1 can1-100 bar1::hisG lys2::HisG pep4::unmarked URA3::pAZ96 (GAL1,10 FLAG-ORC1-opt, ORC2-opt) HIS3::pJF17 (ORC3-opt, ORC4-opt) TRP1::pSG46 (GAL1,10 ORC5-opt, orc6-lnk1scr-C-LPETGG)* | This study |

**Table S3: List of plasmids used in this study**

| Name | Construct | Description | Study |
|---|---|---|---|
| pJF17 | ORC3 + ORC4 | *GAL1,10 ORC3-opt, ORC4-opt* | Frigola et al. 2013 |
| pJF19 | CBP-ORC1 + ORC2 | *GAL1,10 CBP-ORC1-opt, ORC2-opt* | Frigola et al. 2013 |
| pAZ96 | FLAG-ORC1 + ORC2 | *GAL1,10 FLAG-ORC1-opt, ORC2-opt* | Gupta et al. 2021 |
| pSKM002 | MCM4 + MCM5 | *GAL1,10 MCM4, MCM5* | Kang et al. 2014 |
| pSKM003 | MCM6 + MCM7 | *GAL1,10 MCM6, MCM7* | Kang et al. 2014 |
| pSKM004 | MCM2 + FLAG-MCM3 | *GAL1,10 MCM2, Flag-MCM3* | Kang et al. 2014 |
| pSKM033 | FLAG-Cdc6 | *pGEX- GST-PP-FLAG-Cdc6* | Kang et al. 2014 |
| pALS1 | CDT1 + GAL4 | *GAL1,10 CDT1, GAL4* | Kang et al. 2014 |
| pDD48 | ORC5 + ORC6-107-S6 | *GAL1,10 ORC5, ORC6-107-S6* | This study |
| pDD36 | UbSORT-*mcm6Δ103* + MCM7 | *GAL1,10 UbSORT-mcm6Δ103, MCM7* | This study |
| pST048 | MCM2-C-LPETGG + FLAG-MCM3 | *GAL1,10 MCM2C-LPETGG, Flag-MCM3* | Gupta et al. 2021 |
| pAZ87 | ORC5 + UbSORT-*orc6ΔN* | *GAL1,10 ORC5, UbSORT-orc6ΔN* | This study |
| pDD53 | ORC5 + *orc6-107-S6-lnk1scr* | *GAL1,10 ORC5, orc6-107-S6-lnk1scr* | This study |
| pDD49 | ORC5 + UbSORT-*orc6ΔN-lnk1scr* | *GAL1,10 ORC5, UbSORT-orc6ΔN-lnk1scr* | This study |
| pSPB66 | ORC6, URA3 | *ORC6, URA3* | Chen et al. 2011 |
| pRS404-Orc3p-ORC6 | Orc3p-ORC6 | *Orc3pro-ORC6* | Chen et al. 2011 |
| pDD73 | Orc3p-*orc6ΔN* | *Orc3pro-orc6ΔN* | This study |
| pDD74 | Orc3p-*orc6-lnk1scr* | *Orc3pro-orc6-lnk1scr* | This study |
| pDD75 | Orc3p-*orc6ΔN-lnk1scr* | *Orc3pro-orc6ΔN-lnk1scr* | This study |
| pSG26 | Cdt1-C-SORT | *GAL1,10-3xFLAG-Cdt1-C-LPETGG* | Gupta et al. 2021 |
| pRS306 deltN95 Clb5-Flag Cdc28-His | Clb5 + Cdc28 | *GAL1,10 Δ2-95-CLB5-Flag CDC28-His* | Heller et al. 2011 |
| pGEX-Sic1 | Sic1 | *pGEX-Sic1* | Heller et al. 2011 |
| pDD64 | ORC5 + *orc6-107-S6-4A* | *GAL1,10 ORC5, orc6-107-S6-4A* | This study |
| pSG50 | FLAG-ORC1 + *orc2-6A* | *GAL1,10 FLAG-ORC1-opt, orc2-6A* | Amasino et al. 2023 |
| pAZ63 | ORC5 + ORC6-C-LPETGG | *GAL1,10 ORC5-opt, ORC6-C-LPETGG* | Gupta et al. 2021 |
| pSG13 | MCM2 + FLAG-TEV-GG-MCM3 | *GAL1,10 MCM2, Flag-TEV-GG-MCM3* | Gupta et al. 2021 |
| pSG46 | ORC5 + *orc6-lnk1scr*-C-LPETGG | *GAL1,10 ORC5-opt, orc6-lnk1scr-C-LPETGG* | This study |
